# A dual readout embryonic zebrafish xenograft model of rhabdomyosarcoma to assess clinically relevant multi-receptor tyrosine kinase inhibitors

**DOI:** 10.1101/2024.12.19.629341

**Authors:** Joseph W. Wragg, Emma L. Gray, Rui Monteiro, Jo R. Morris, Andrew D. Beggs, Ferenc Müller, Susanne A. Gatz

## Abstract

**Background:** Rhabdomyosarcoma (RMS) is a highly aggressive soft tissue sarcoma, affecting children and adolescents, with poor prognosis in some patient groups. Better therapeutic regimens and preclinical models to test them in are needed. Multi-receptor tyrosine kinase inhibitors (MRTKIs) are licensed for adult indications and explored in the clinic in sarcoma patients. The MRTKI Regorafenib is currently assessed in the relapse setting in patients with RMS (NCT04625907). Reliable biomarkers of response for MRTKIs are lacking. MRTKIs act not only against the cancer cell, but also the supporting stroma, particularly the vasculature. The embryonic zebrafish is translucent and allows assessment of this interaction with high-throughput *in vivo* imaging.

**Methods:** A new preclinical embryo zebrafish xenograft model was developed using Tg(*flk1*:GFP) (blood vessel reporter) transgenic zebrafish embryos inoculated in the yolk with fluorescently labelled cells from 7 different RMS cell lines (fusion-positive (FP): Rh4, Rh30, Rh41, RMS-01, fusion-negative (FN): RD, JR1, SMS-CTR), and patient-derived cells IC-pPDX-104 at 50 hours post-fertilization and incubated at 34°C for up to 70 hours. Xenografts and vessel beds were imaged and analysed using custom FIJI pipelines. MRTKIs regorafenib and infigratinib were used at a concentration of 0.1uM added to the fish water 4 hours post cell inoculation. Pro-angiogenic growth factors VEFG-A, FGF-2 and PDGF-BB were measured in conditioned media of each cell line.

**Results:** All 7 RMS cell lines and the patient-derived cells engrafted with tumour burden assessment by fluorescent imaging and direct cell counting indicating adequate growth and high cell viability during the observation period. RMS tumours induced neo-vascularisation towards the tumour and increased density of proximal vessel beds. MRTKI treatment revealed a greater tumour-intrinsic sensitivity of FP cells, but identified a significant blockade of neo-vascularisation across all RMS lines, with regorafenib response correlated with secretion of VEGF-A.

**Conclusion:** We have developed an embryonic zebrafish xenograft model of RMS, which allows assessment of tumour growth, vascularisation initiation and therapeutic responses to clinically relevant MRTKIs. The identification of VEGF-A secretion as potential biomarker for Regorafenib response and the separation of therapeutic effects on tumour growth and neovascularisation suggests additional value of our model for response prediction to MRTKIs.

## Introduction

Rhabdomyosarcoma (RMS) is a highly aggressive soft tissue sarcoma, predominantly affecting children and adolescents. It constitutes 3-4% of all childhood malignancies and 50% of childhood soft tissue sarcomas. This represents around 350 cases per year in the US, with similar incidences in the UK and Europe (1, 2). Treatment includes intensive chemotherapy, surgery and radiotherapy and outcome has remained largely unchanged over the last decades with 5-year survival still well below 30% for some cohorts of patients (2–4). In addition, even when classified as disease-free, survivors often experience long term morbidities owing to the use of aggressive multi-modal treatment (3, 4).

The majority of childhood and adolescent RMS fall into two histological subtypes, embryonal (ERMS) and alveolar (ARMS). These two subtypes also broadly subdivide by molecular drivers, with 80% of ARMS associated with chromosomal translocations t(2;13)(q35;q14) or t(1;13)(p36;q14), leading to the formation of a fusion of *PAX3* or *PAX7* with *FOXO1* (1, 2, 5). This PAX3/7-FOXO1 fusion protein constitutes a key driver of malignancy in so-called Fusion-positive (FP)-RMS, establishing an aberrant myogenic super-enhancer program and transcriptionally activating oncogenic targets including the *FGFR* and *PDGFR* gene families of receptor tyrosine kinases, alongside *ALK* and *MYC* genes (5–9). ERMS are more genetically diverse, lacking the PAX3/7-FOXO1 fusion protein, but frequently harbour mutations in the MAPK, RAS and phosphatidylinositol-3-kinase (PI3K) pathways, which also lead to the aberrant activation of upstream receptor tyrosine kinases (1, 5, 8, 10). ARMS lacking the PAX3/7-FOXO1 fusion protein are molecularly and clinically indistinguishable from ERMS, together constitute the fusion-negative (FN)-RMS group (1, 5, 11), and have a better outcome, whereas FP-RMS is associated with significantly worse outcomes which led to current prospective clinical studies using fusion status rather than histology as risk stratifier (2, 12, 13).

Intriguingly the differing molecular events defining FP-RMS and FN-RMS both converge on the over-activation of receptor tyrosine kinase signalling through the MAPK and PI3K pathways (1, 8). This shared molecular driver represents a potential target for therapeutic intervention that would benefit a wide range of RMS patients. Therefore, an emerging therapeutic angle in RMS is the use of multi-receptor tyrosine kinase inhibitors (MRTKIs), an important class of agents, many of which are licensed for adult indications and explored in the clinic in sarcoma patients including children and young adults (14, 15). The MRTKI regorafenib is a potent inhibitor of VEGFR1-3, but also PDGFR, FGFR1/2 amongst others (15). In pre-clinical testing, it has shown moderate growth inhibition against RMS cell lines in vitro and significant growth delay *in vivo*, albeit with only one line tested (16). Following on from that, regorafenib was assessed in a Phase I as single agent in recurrent or refractory paediatric solid tumours and in a subsequent Phase 1b in combination with backbone chemotherapy vincristine and irinotecan (VI) in recurrent or refractory paediatric solid tumours with focus on RMS (both NCT02085148). Two of three RMS patients had some clinical benefit with the single agent therapy (one unconfirmed partial response, one prolonged stable disease) (17) and seven out of 12 RMS patients exhibiting either a complete response (n=2) or partial response (n=5) (18). Given the promising results in RMS patients, the VI + regorafenib combination was introduced into the current European platform trial “frontline and relapsed – RMS” (FaR-RMS, NCT04625907) in the relapse setting and randomised against the established standard of care of VI + temozolomide (2, 4). Another MRTKi, infigratinib (a potent inhibitor of FGFR1-3) has also shown promising results in pre-clinical testing *in vitro* and *in vivo*, particularly against FP-RMS, but clinical testing has so far been restricted to adult cancer types (19, 20). Whilst detection of *FGFR2* gene fusions is an accepted pre-selection biomarker in cholangiocarcinoma patients for infigratinib therapy (21), biomarkers of response for regorafenib or infigratinib in the clinical setting in sarcoma including RMS are currently unclear (15).

Growth factor signalling through the targets of both drugs, regorafenib and infigratinib, are important for not only the cancer cell, but also the supporting stroma, particularly the vasculature. The interaction of the tumour with its vasculature and the effect of MRTKIs on this cannot be modelled in standard cell culture and investigation of these interactions and effects is difficult to assess in *in vivo* mouse xenograft studies, as it cannot be directly visualised due to the opacity of the model. Hence, current cancer drug development methods are insufficient for a comprehensive assessment of MRTKIs and the development of biomarkers for response.

The embryonic zebrafish (<120 hours post fertilisation [hpf]) is close to transparent allowing detailed visualisation of internal structures such as the vasculature identifiable with the aid of *flk-1* or *fli-1* transgenic reporters and the use of high throughput *in vivo* imaging (22–25). Additionally, the ability of this model system to in principle assess the effect of anti-angiogenic therapeutics has previously been demonstrated in lung, breast and colorectal carcinoma models, using mostly one to two and a maximum of three (colon) cell lines per disease type (26–29). Overall, the embryonic zebrafish is emerging as a promising host organism for xenograft tumour modelling and assessment of stromal responses (30–32).

We, here, share the development and detailed analysis of a new embryonic zebrafish model for assessment of RMS cell lines and patient derived tumour cells including their effect on the host vasculature and the response of the injected tumour xenografts and host vasculature to treatment with the clinically relevant MRTKIs, regorafenib and infigratinib. This model system reveals potential biomarkers for future testing and is expected to add additional value to the standard preclinical assessment tools of 2D/3D cell culture analysis and *in vivo* xenograft mouse models. Further parallel assessment of patient-derived tumour cells in our new model system alongside conventional preclinical studies and, ultimately, the use within a co-clinical trial setting will establish its full value.

## Materials and Methods

### Zebrafish handling

All animal husbandry and associated procedures were approved by the British Home Office (Licence number: PP2470547). Zebrafish embryos were obtained by sibling crosses from adult *Tg(flk1:GFP)* fish housed in the University of Birmingham fish facility. Zebrafish were bred and embryos raised and staged following standard protocols (33, 34).

### Cell culture

RD, JR1, SMS-CTR, Rh30, Rh4, Rh41 and RMS01 cell lines were kindly provided by Professor Janet M. Shipley (details of cell origins are as in (19)). IC-pPDX-104 cells were kindly provided by Professor Beat Schäfer with use covered by MTA2017008 between University Hospital Zurich and University of Birmingham. IC-pPDX-104 were originally generated within the laboratory of Professor Olivier Delattre at the INSERM-Institut Curie. RD, Rh30, Rh4, SMS-CTR, RMS01 and HFF-1 (ATCC) were cultured in Dulbecco’s Modified Eagle Medium (Gibco), JR1 and Rh41 cells were cultured in RPMI-1640 medium (Gibco). Each were supplemented with 10% foetal bovine serum (Gibco), 1% penicillin/streptomycin (Gibco) and 2mM Glutamax (Gibco). IC-pPDX-104 cells were cultured on Matrigel (Corning) coated plates in Advanced DMEM (Gibco), supplemented with B27 (Life Technologies), 1.25mM N-acetyl-L-cysteine (Sigma-Aldrich), 20 ng/ml bFGF (PeproTech), 20 ng/ml EGF (PeproTech) 1% penicillin/streptomycin (Gibco) and 2mM Glutamax (Gibco) (35). All cell lines were maintained in a tissue culture incubator at 37°C in 5% carbon dioxide. Identity of all cell lines was confirmed by STR testing (Eurofins). Cells were regularly tested for mycoplasma by PCR following established protocols (36).

### Cell preparation for microinjection

Culture media were removed from cells grown to ∼80% confluence. The cells were then washed with phosphate-buffered saline (PBS) and Trypsin (Gibco) was added to release them from the culture plate. The Trypsin was neutralised through addition of 7x volume culture media. Cells were then collected, centrifuged at 300g for 5 minutes and washed with Optimem (Gibco), repeating the centrifugation supernatant removal step. Cells were re-suspended at a density of 1×10^6^ cells/ml in cell staining solution (Vybrant CM-Dil cell labelling solution (Invitrogen) diluted 1:250 in Optimem [Gibco]) and incubated for 20 mins at 37°C in a tissue culture incubator, followed by 5 mins on ice. Cells were washed twice by centrifugation at 4°C and resuspension in ice cold PBS. Cells were finally re-suspended at a density of 2.5×10^5^ /µl in microinjection solution (0.5mg/ml Collagen I [Rat Tail, Gibco] in PBS) and kept on ice prior to injection.

### Zebrafish embryo microinjection

*Tg(flk1:GFP)* embryos were maintained to 50 hours post fertilisation (hpf) [long pec] at 28°C in E3 medium. They were then decorionated with 1mg/mL Pronase and anaesthetised immediately prior to microinjection using 0.2mg/ml Tricaine (Ethyl 3-aminobenzoate methanesulfonate) solution (Sigma-Aldrich). The injection solution (containing either cells as previously described, or recombinant growth factors suspended at the appropriate concentration in PBS, as well as 1:10 Phenol red solution [Gibco] to aid injection site determination) was back-loaded into borosilicate glass microcapillary injection needles (Warner Instruments) and mounted onto the micro-injector (Tritech). Anaesthetised embryos were aligned and injected with 1nl of injection solution into the centre of the yolk sac.

### Microinjected zebrafish embryo maintenance and drug treatment

Following microinjection, the embryos were transferred into fresh E3 medium, transferred to a 28°C incubator for 30 mins, and then to a 34°C incubator for 3.5 hours. 54 hpf Embryos were then screened under the fluorescent microscope for consistency in injection mass placement and size, with non-compliant embryos discarded. Zebrafish embryos microinjected with tumour cells or growth factors were then randomly assigned into drug treatment groups and systemically treated with 0.1μM regorafenib (SelleckChem), 0.1μM infigratinib (SelleckChem), or 1:100.000 DMSO dissolved in E3 medium. During the assay development phase also higher and lower concentrations of drugs were used. Embryos were then maintained at 34°C until 120 hpf (Early larvae).

### Zebrafish embryo imaging and analysis

Microinjected zebrafish embryos were retrieved from the 34°C incubator at 120 hpf, anaesthetised with 0.2mg/ml Tricaine/ E3 solution and transferred to a ZF 96 well imaging plate (Hashimoto). Embryos were aligned under a stereo microscope and the plate imaged using Cytation 5 microscope (Agilent), with Gen5 software (Agilent) calibrated for imaging. Regions of interest (ROI) were manually selected for each well to centre the sub-intestinal vessel region, and embryos were imaged at 10X magnification on the GFP, RFP and brightfield channels, as appropriate. The images were then processed into Z-stacks within Gen5, and exported as TIF files for quantification and analysis. Following imaging, the embryos were euthanized by a schedule 1 method.

### Image Quantification and Analysis

Image quantification and analysis was performed in Fiji. TIF files were loaded into Fiji. Tumour area and fluorescence was determined using the following custom macro:

*run(“8-bit”);*

*setThreshold*(*173, 255*);

*run(“Convert to Mask”);*

*run(“Set Measurements…”, “area integrated area_fraction redirect=None decimal=3”);*

*run(“Measure”);*

For vessel length measurement the ROIs for neo-vessels, Sub-intestinal vessels (SIV) and intersegmental vessels (ISV) were defined as described in Figure 2A and the “Clear outside” command used to remove the rest of the image. The following custom macro, adapted from (37) was then used to quantify each vessel within the ROI:

*run(“Subtract Background…”, “rolling=50”);*

*run(“Enhance Contrast…”, “saturated=0.3”);*

*run(“Despeckle”);*

*run(“Remove Outliers…”, “radius=2 threshold=50 which=Bright”);*

*run(“Tubeness”, “sigma=2 use”);*

*run(“Enhance Contrast…”, “saturated=0.10”);*

*setOption(“ScaleConversions”, true);*

*run(“8-bit”);*

*setOption(“BlackBackground”, false);*

*run(“Convert to Mask”);*

*run(“Invert LUT”);*

*run(“Skeletonize (2D/3D)”);*

*run(“Analyze Skeleton (2D/3D)”, “prune=none show display”)*

This macro utilises the subtract background, enhance contrast and tubeness filters in Fiji to pre-process the images to enhance filamentous vessels. The images are then thresholded to create binary images, before the ‘Skeletonize’ plugin reduces the segmented vessels to single-pixel-wide lines. The ‘Analyze Skeleton’ plugin then provides the length of each vessel in pixels, which was summed to calculate the total vessel length for the parameter of interest.

**Figure 1:**
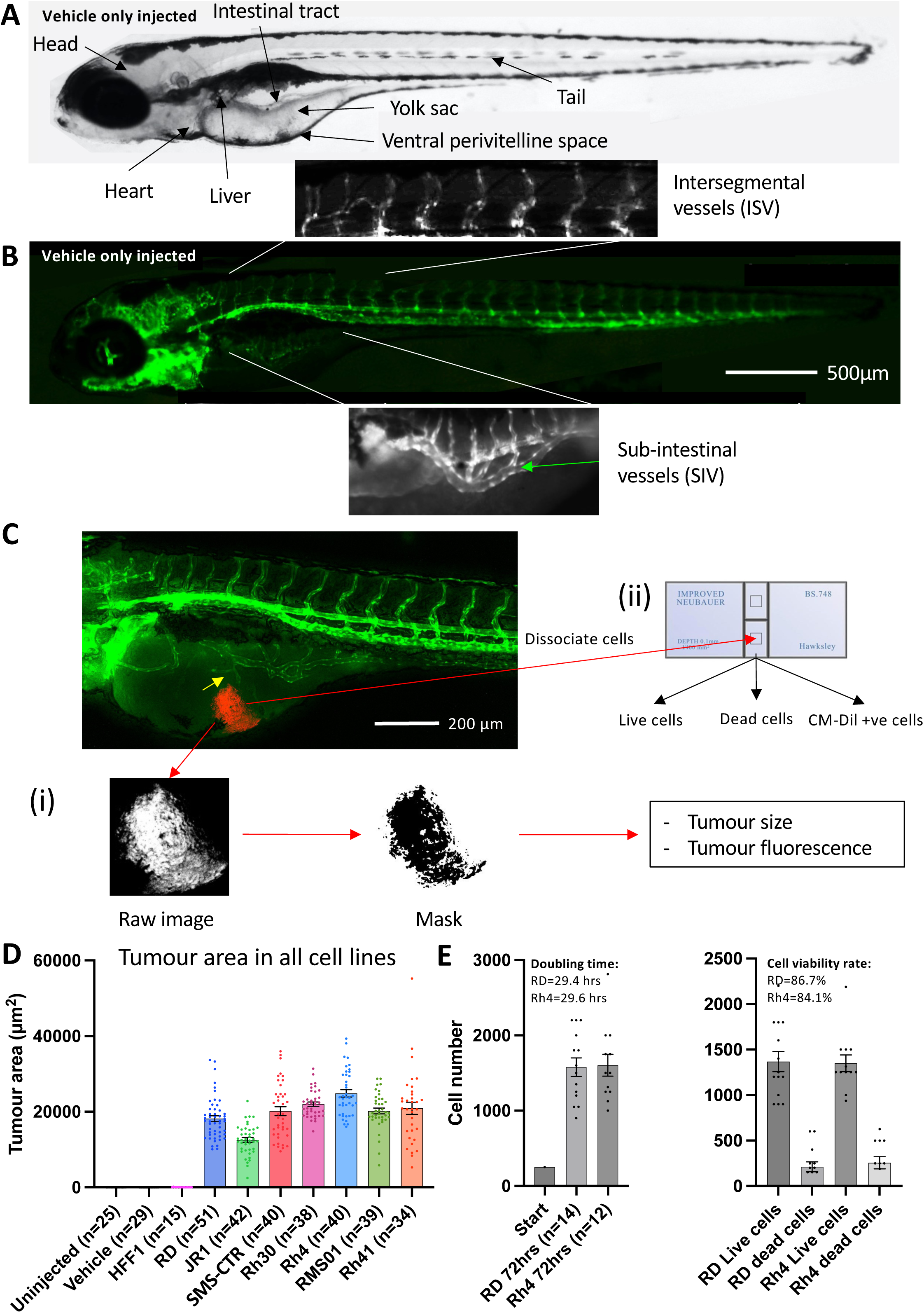
RMS tumour cells are viable and proliferate in larval zebrafish hosts. **A**, Brightfield image of a 120 hpf zebrafish embryo highlighting key anatomical features, relevant to xenograft creation. **B**, Matched fluorescent image illustrating the marking of developing blood vessels by the *flk1:GFP* transgene (green). The sub-intestinal vessel (SIV) and intersegmental vessel (ISV) beds are highlighted. **C**, Schematic illustrating the techniques used to measure tumour burden. An image of a 120 hpf (70 hpi) zebrafish xenograft in *flk1-GFP* background is shown with blood vessels shown in green and the RMS tumour in red. Tumour induced reconfiguration of the SIV forming neo-vessel sprouts is marked (yellow arrow). **(i)** tumour size and fluorescence are assessed through FIJI image processing and analysis and **(ii)** tumour cell number and viability are assessed by excision of the tumour and dissociation, followed by trypan blue staining and cell counting on a haemocytometer. **D**, Tumour area was calculated from fluorescent images as described in C, for 7 RMS cell lines alongside human foreskin fibroblasts (HFF1) as well as un-injected and vehicle (collagen) only injected zebrafish embryos. Data are presented as mean values +/− SEM. Data from 5 independent experiments for each line, with the exception of RD and HFF1 where n=6 and 3, respectively. **E**, Bar chart showing total (left) and live/dead (right) cell number counts from excised RD and Rh4 tumours, measured at the end of the xenograft experiment (70 hpi) and compared to the initial injected bolus of 250 cells. Doubling time and cell viability rate are shown. Data from 2 independent experiments. n = number of xenografts.

**Figure 2:**
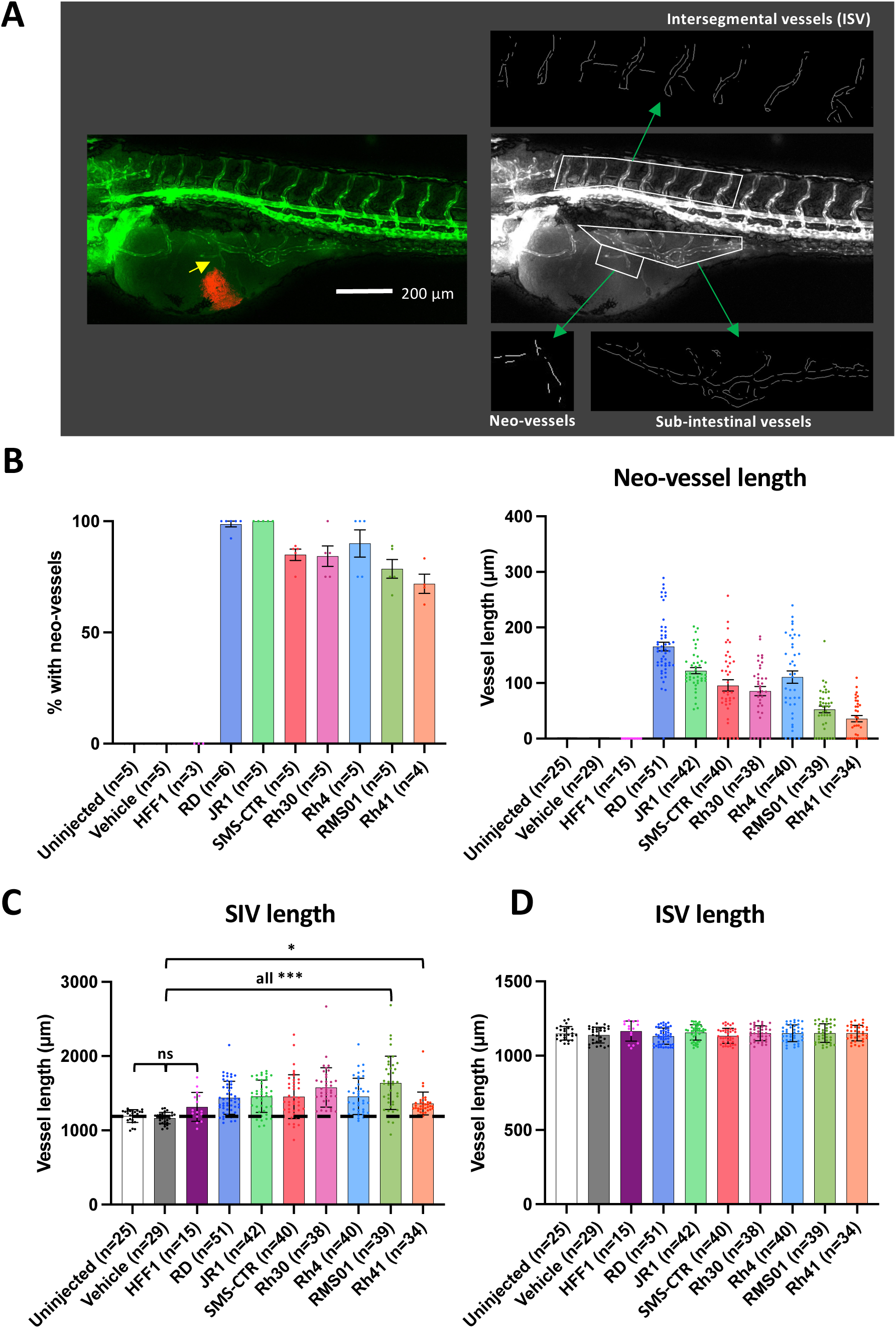
RMS tumours induce neo-vascularisation from proximal vessel beds. **A**, Schematic illustrating the calculation of sub-intestinal, intersegmental and tumour induced neo-vessel lengths, by Fiji image analysis. An image of a 120 hpf (70 hpi) zebrafish xenograft in *flk1-GFP* background is shown with blood vessels shown in green and the RMS tumour in red. From this, the green fluorescence channel is taken, GFP +ve vessels skeletonised and their lengths measured. **B**, Bar graphs showing the proportion of xenografts with neo-vessel induction for xenografts generated from each RMS cell line (left) and the average neo-vessel length for xenografts generated from each RMS cell line (right). **C** and **D**, Bar graphs showing the average sub-intestinal vessel length (excluding neo-vessels) (**C**) and the average intersegmental vessel length (**D**) for xenografts generated from each RMS cell line. Dotted line indicates level of vehicle only injected zebrafish controls. * P<0.05, *** P<0.001, Ordinary one-way ANOVA with Tukey’s multiple comparison’s test, Data are presented as mean values +/− SEM. n = number of xenografts (except for B left, which is number of experiments).

### Cell culture drug response assessment, imaging and analysis

5000 cells from each cell line under investigation were seeded in 96 well plates in the appropriate media and culture conditions described earlier. Cells were allowed to settle for 24hrs prior to the media being removed and replaced with media containing the appropriate concentration of regorafenib or infigratinib or solvent (DMSO). Cells were imaged daily using the Brightfield high contrast lens on the Cytation 5 microscope (Agilent) for 7 days, with a media change after 3 days. Confluence analysis was performed on the cell images using Gen5 software (Agilent), with the following settings: Threshold: 3500, Background: Light, Split touching objects, Fill holes in masks, Admissible object size: 30-3000. Comparative confluence analysis for treatment response was performed for each cell line at the time-point where the control group reached 100% confluence (typically 5-7 days). GI50 (50% inhibition of maximum confluence) was quantified from this data using Graphpad PRISM.

### Enzyme-Linked Immunosorbent Assay (ELISA)

250,000 cells from each cell line were plated in 6 well plates and cultured for 24hrs. The culture media was then replaced with DMEM minus FBS (DMEM supplemented with Glutamax [Gibco] and Pen/Strep [Gibco]) for 24 hours prior to media harvesting. DMEM minus FBS incubated alongside without cells was additionally tested as a control. The media were then tested on the Human Angiogenesis ELISA Profiling Assay kits I&II Chemiluminescence (Signosis) following manufacturer’s protocols. In brief, 100μL of media to be tested was added to each well of the ELISA plate and incubated for 2 hours at room temperature with gentle shaking. The wells were then aspirated and washed 3 times with 200μL wash buffer. Next, 100μL of biotin-labelled antibody mixture was added to each well and incubated for 1 hour at room temperature with gentle shaking. The wells were then washed 3 times with 200μL wash buffer. 100μL of streptavidin-HRP conjugate was added to each well and incubated for 45 minutes at room temperature with gentle shaking. The wells were then washed a further 3 times with 200μL wash buffer, with the wash buffer incubated in the wells for 10 minutes during each wash in order to reduce background signal. Finally, 95μL of substrate solution was incubated for 2 minutes in each well, and the plate was then immediately read in a luminometer. The integration time was set to 1 second with no filter position, and each well was read in triplicate. The triplicate counts were then averaged and normalised to the average readings from the cell free control wells. Absolute concentration of secreted growth factors was determined by addition of serially diluted protein standards (Signosis) to the DMEM minus FBS control media to a final concentration of 8, 4, 2, 1, 0.5 ng/ml of each growth factor, which was then tested on the ELISA plate. From this, standard curves were generated to determine absolute concentration of secreted growth factors.

### Ethics statement

This research uses no fresh human material, or cells directly from patients. All experiments are performed with RMS cell lines and, in the case of IC-pPDX-104, a short-term *in vitro* culture derived from a patient derived xenograft which had been developed through serial passaging in mice. Adult zebrafish maintenance and all embryonic zebrafish work was carried out under the British Home Office Licence number: PP2470547.

### Statistics

All statistical analysis was performed using the Graphpad Prism software package (version 9, Boston, MA, USA). All data figures were also generated using the Graphpad Prism software package. Data are presented as mean values +/− SEM unless otherwise stated. Experimental replicates and n-numbers are also stated in the figure legends.

## Results

### RMS tumour cells are viable and proliferate in embryonic zebrafish hosts

We aimed to generate an embryonic zebrafish xenograft model of rhabdomyosarcoma, allowing assessment of tumour behaviour and stromal interactions; i.e., change in host vessels and neo-vessel induction, without treatment or in the presence of therapeutics. For this purpose we selected the *Tg(flk1:GFP)* zebrafish line, widely used for profiling of zebrafish vessel development (22–25, 38) (Figure 1A&B), as xenograft host for RMS cells fluorescently labelled with CM-Dil, to allow assessment of tumour burden (Figure 1C, Supplementary figure 1A).

We first looked to identify the optimal site for tumour cell microinjection, assessing the two most commonly used injection sites, the yolk and perivitelline space (PVS) (Figure 1A, Supplementary figure 1B (i)). We injected 250 cells from two FN-RMS (RD and SMS-CTR) and two FP-RMS lines (Rh30 and Rh4), into both sites of 50 hpf (High Pec) zebrafish embryos, in 3 matched experimental rounds. Seventy hrs after RMS tumour inoculation the xenograft model was imaged by fluorescent microscopy to assess tumour area and fluorescence. This was then quantified using a custom Fiji pipeline, that generated a fluorescence signal informed mask, from which tumour area and fluorescence could be quantified (Figure 1C (i)). Intriguingly, this assessment revealed that inoculation into the yolk formed on average 10-fold larger tumours than injection into the PVS for all lines (2310 vs. 22919 μm^2^) (Supplementary figure 1B (ii)). In addition to this, between 75-96% of yolk-inoculated xenografts induced the development of neo-vessels from the proximal sub-intestinal vessel (SIV) bed, compared to 3.4-14.8% for PVS inoculated xenografts (Figure 1C, Supplementary figure 1B (iii)). No alternative source of co-opted vessels was observed in the PVS-inoculated xenografts. As both, tumour size and vessel induction, were key metrics which we wanted to assess with this model system, we decided to use the yolk as the injection site of choice.

We next looked to assess whether a broader range of RMS tumour cell lines would establish tumours in zebrafish embryonic hosts and made sure to include cell lines which had previously been used in *in vivo* experiments in mice when addressing response to MRTKIs for later comparison (regorafenib: RMS01 (16); infigratinib: Rh41 and RMS01 (19)). We injected embryos with a total of three FN-RMS lines (RD, JR1 and SMS-CTR) and four FP-RMS lines (Rh30, Rh4, RMS01 and Rh41), alongside human foreskin fibroblast (HFF1) cells as a non-cancerous control and quantified the tumour area based on fluorescence. This assessment revealed that each of the RMS lines establish tumours in zebrafish embryos, whereas the non-cancerous HFF1 cells do not engraft (Figure 1D).

For implanted RMS tumours to accurately recapitulate clinical behaviour they must be viable and able to proliferate, we therefore assessed the cell doubling time and viability rate of xenografts formed from two RMS cell lines (RD and Rh4), representing one FN-RMS and FP-RMS, respectively. In addition to the fluorescent tumour size assessment, tumours were additionally excised from the xenografts by microdissection of the yolk region to assess cell number and viability by trypan blue exclusion assay (Figure 1C (ii)). Using these quantifications, we compared the total RMS cell number after 70hrs in the xenograft (RD – 1578, Rh4 – 1604) to the initial inoculation cell number (250 cells), providing a doubling time for each of the lines within the xenograft model system (RD – 29.4hrs, Rh4 – 29.6hrs) (Figure 1E). These findings reveal that both classes of RMS tumours are growing in the xenograft model system. To assess cell viability, we further separated the total cell number in live and dead cells as assessed by presence/absence of trypan blue intracellular staining, respectively. This analysis revealed a cell viability rate of 86.7 and 84.1% for RD and Rh4 tumour cells, respectively (Figure 1E).

To ensure that the cells excised from the yolk for counting were the fluorescently labelled RMS cell lines, we repeated the cell count, doubling time and viability assessments on CM-Dil positive cells only. This analysis revealed that live cells and CM-Dil positive cell numbers are very closely aligned, but that very few dead cells were CM-Dil positive (Supplementary figure 1C). This could suggest that the majority of non-RMS cells in the tumour are dead, but the more likely explanation is that RMS cells lose their fluorescence upon apoptosis, suggesting that a fluorescence-based assessment of tumour burden could be used as a robust proxy for live tumour cell number. To test this, we measured tumour area and fluorescence from fluorescent images matched to the xenografts used for cell counting (Supplementary figure 1D) and performed Pearson correlation analysis between live, dead, CM-Dil positive and total cell numbers with tumour area and fluorescence as quantified by fluorescence imaging and Fiji pipeline quantification. This analysis revealed a strong correlation between both tumour area and fluorescence measurements with live cell number (r=0.88 and 0.86 respectively) with a weaker correlation with total cell number and anti-correlation with dead cell number (Supplementary figure 1E). This is encouraging as it suggests that tumour area and fluorescence measurements are reflective of the amount of viable tumour in the xenograft. As tumour area quantification showed the highest level of correlation with live cell count, we decided to use this metric for quantification of RMS xenografted tumours moving forward. It should be noted that the tumour is a 3-dimensional object which we measure here in 2D terms due to difficulty in extrapolating the volume of amorphous shaped tumours. Therefore, the measurements of changes in tumour size may be an underestimation but are likely proportional to true volumetric change.

### RMS tumours induce neo-vascularisation from proximal vessel beds

Alongside RMS tumour establishment and growth, we also wanted to assess the interaction of RMS tumours with the host blood vessels. As previously noted RMS xenografts inoculated in the yolk consistently induced neo-vessel sprouting from the proximal SIV bed (Figure 1C, Supplementary figure 1B (iii)). We reassessed this for the full range of RMS xenografts described in Figure 1D, finding that neo-vessel induction efficiency ranged from 71.5% in Rh41 to 100% in JR1 xenografts (Figure 2A and B). This indicates that RMS tumours are capable of inducing angiogenesis and acquiring vascularisation in this model system. In order to quantitatively assess the extent of induced neo-vessel production, as well as the effect of each of the 7 RMS tumour lines on proximal (SIV) and distal (intersegmental vessel [ISV]) vessel beds we generated a custom Fiji pipeline (37) to skeletonise and measure these three vessel populations from fluorescent images of the xenograft model system (Figure 2A). This analysis revealed that each of the RMS tumour lines induces both, neo-vessels (Figure 2B) and a significant increase in SIV length (Figure 2C), whereas the ISV length remained unaffected (Figure 2D). This is consistent with RMS tumours inducing neo-vessel sprouts and stimulating increased vessel complexity in existing proximal beds and suggests this effect is spatially constrained to the proximity of the tumour.

Intriguingly, FN-RMS lines appear to be more efficient at inducing neo-vessels (average % with neo-vessels: 94.5% FN-RMS vs. 81.1% FP-RMS; average neo-vessel length: 127.9 µm FN-RMS vs. 71.1 µm FP-RMS), whereas FP-RMS stimulate SIV development with greater efficiency (% change in SIV length versus vehicle injected: 24.5% FN-RMS vs. 29.1% FP-RMS) (Supplementary figure 2A).

The variation in neo-vessel length between xenografts could be a consequence of either varying neo-vessel induction or elongation efficiencies (or both). As these reflect distinct stages of angiogenesis the efficiency of which may vary between xenografts, we repeated the neo-vessel length analysis excluding xenografts where no neo-vessels had been induced, ie focusing on elongation efficiency only. This analysis again revealed FN-RMS to have the greater neo-vessel length (average neo-vessel length upon induction: 134.6 µm FN-RMS vs. 85.4 µm FP-RMS) (Supplementary figure 2B).

### Identification of a suitable dose of MRTKI regorafenib and infigratinib in non-tumour bearing embryonic zebrafish for subsequent xenograft exposure

As we have established a model system where RMS tumours grow and vascularise robustly, we next sought to identify whether it could be used to investigate the effect of the clinically relevant MRTKIs regorafenib and infigratinib on RMS tumour growth and angiogenesis.

Regorafenib is a potent inhibitor of several different receptor tyrosine kinases with relevant targets in cancer on both, blood vessels and stromal cells (amongst them VEGFRs, PDGFRs and FGFRs) (15), whereas infigratinib is a potent, selective inhibitor of FGFR1-3, and >10-fold lower sensitivity towards FGFR4 found on both tumour and endothelial cells (19, 39).

Therefore, as next step in the development of our model, we needed to establish whether the embryonic zebrafish host, which provides the tumour stroma including the endothelial cells, is responsive to MRTKI blockade through regorafenib and infigratinib, outside of the tumour context. To this end, we assessed the effect of various drug concentrations of regorafenib and infigratinib on zebrafish embryonic development to identify the maximum dose of these drugs at which no obvious impact on development – with particular emphasis on vascular development – could be identified. Any drug effects observed in the xenograft model applying this new “no adverse effect drug dose” will be due to impacts on the tumour and associated vasculature. To achieve this, we treated tumour-free 50 hpf (High Pec) zebrafish embryos with serial dilutions of each drug between 0.05 and 10 µM supplied in the fish water for 66 hours (mirroring the tumour growth period) and measured ISV and SIV lengths (Supplementary figure 3A and B) as previously described, as well as assessed effects on embryonic development (Supplementary figure 3C and D). This analysis identified no significant impact of regorafenib on either vessel bed length or embryonic development at a concentration of ≤0.1 µM. Higher concentrations displayed, both, an induction of pericardial oedema and a significant reduction in SIV and ISV lengths. This is consistent with a previous study investigating regorafenib toxicity in zebrafish (40). Infigratinib had no significant effect on normal vessel development at a concentration of ≤0.5 µM, with overall development not affected at a concentration of ≤0.1 µM (Supplementary figure 3). Like regorafenib, higher concentrations of infigratinib disrupted normal embryonic vascularisation (although to a much lesser extent) and induced pericardial oedema, but additionally induced tail curvature at concentrations of ≥0.5 µM. We, therefore, decided to use a drug concentration of 0.1 µM also for infigratinib in future xenograft experiments.

### MRTKIs regorafenib and infigratinib block growth of RMS xenografted tumours

We next treated RMS xenografts generated from the 7 cell lines, with 0.1 µM regorafenib or infigratinib and assessed the effect on tumour area. Strikingly, we found that both drugs caused significant reduction in tumour area for all RMS lines (Supplementary figure 4). The extent of tumour growth reduction induced by the MRTKIs was however heterogeneous between the lines (Figure 3A). With both, regorafenib and infigratinib, having a greater effect on FP-RMS tumours than FN-RMS tumours, and infigratinib being particularly potent against FP-RMS tumours (average relative tumour area; 0.52 vs. 0.41 regorafenib and 0.52 vs 0.29 infigratinib, for FN– and FP-RMS respectively), although not statistically significant for either drug (P=0.19 and 0.08, respectively; two-tailed T-test). This is likely due to the results for the relatively resistant FP-RMS line Rh30 and was previously observed *in vitro* for both drugs (16, 19) (Figure 3A). To further validate these findings, we performed tumour area measurements and direct cell counting analysis on a subset of RD and Rh4 xenografts as previously described (Supplementary figure 5). In agreement with the overall findings, this analysis also showed a significant reduction in tumour cell viability for both drugs, with the greatest potency displayed by infigratinib in the FP-RMS line Rh4 (Supplementary figure 5). Interestingly, infigratinib treatment showed different effects in the two cell lines. In Rh4 the effect of the drug was primarily on cell viability, whereas in RD the observed reduction in tumour area appeared to be the result of reductions in both viability and proliferation rate (Supplementary figure 5).

**Figure 3:**
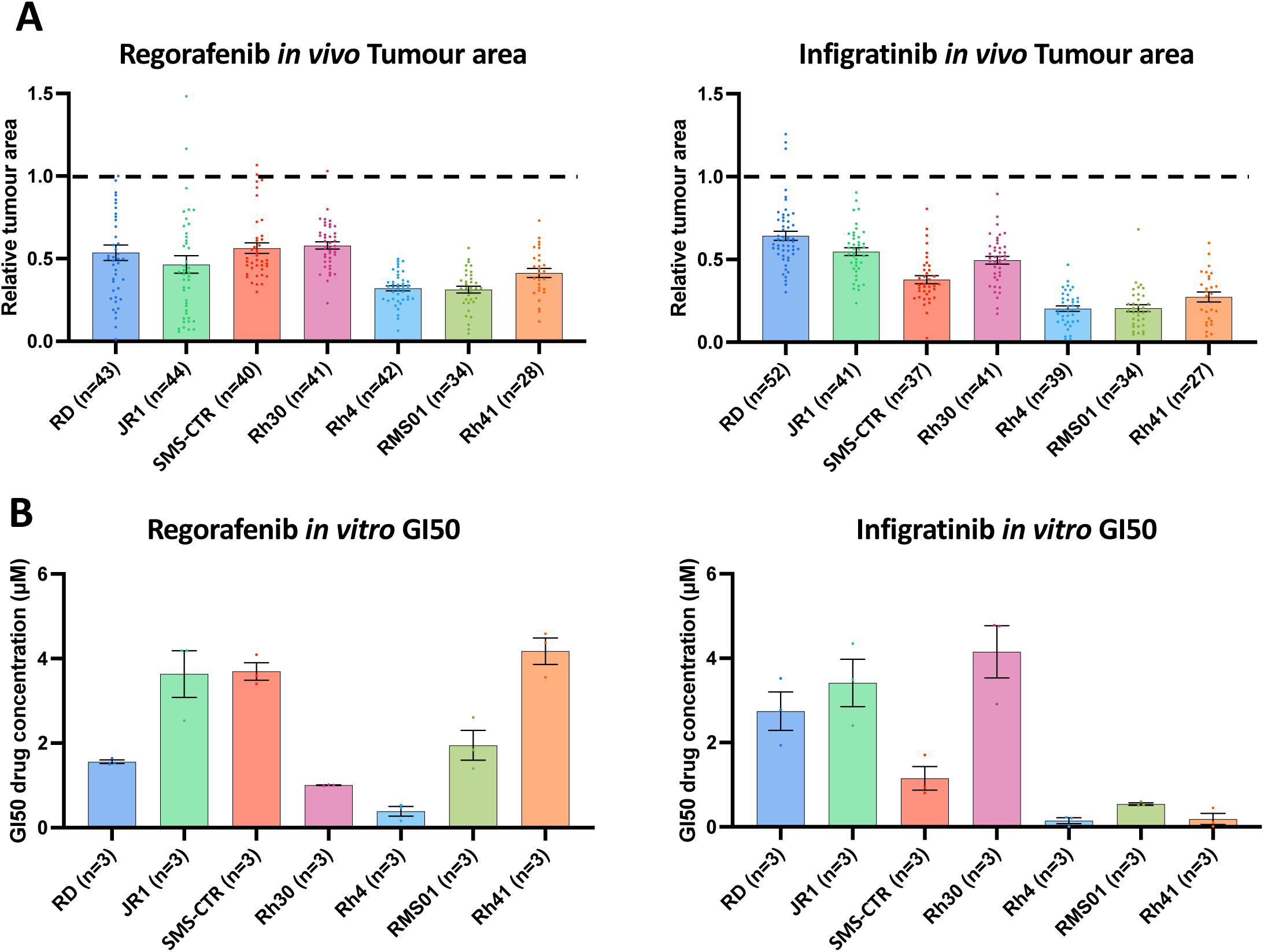
Regorafenib and infigratinib treatment inhibits RMS xenograft tumour growth. **A**, Xenografts generated from the 7 RMS cell lines were treated with 0.1 µM regorafenib and infigratinib for 66 hours. Bar graphs of tumour area for treated xenografts relative to the average vehicle treated tumour area for each cell line. Dotted line indicates level of relative tumour area for vehicle treated xenografts (1.0 =100% for each cell line). B, RMS cell lines were treated *in vitro* in 2D with regorafenib and infigratinib at a range of concentrations (0.1-10 µM) for 7 days (as described in Supplementary figure 6a) with the drug concentration inducing a 50% reduction in maximum cell surface, relative to vehicle treated (GI50) calculated and shown in the bar graphs. ** P<0.01, **** P<0.0001, Ordinary one-way ANOVA with Tukey’s multiple comparison’s test, Data are presented as mean values +/− SEM. n = number of xenografts

To further validate these findings, and additionally to compare the pattern of RMS tumour response to regorafenib *in vitro* versus our *in vivo* model system, we performed an *in vitro* assessment of regorafenib and infigratinib drug response for each of the 7 RMS lines. We treated the lines with serial dilutions of 0.1-10 µM for both drugs, identifying the GI50 concentration for each line using confluency as readout (Figure 3B, Supplementary figure 6). This analysis revealed for infigratinib a striking level of consistency in relative responsiveness of RMS tumour *in vivo* and cell confluency *in vitro*, respectively, between the lines, whereas, for regorafenib, relative responsiveness was quite different between the model systems (Figure 3A and B). Intriguingly, however, RMS tumours appear to be universally more sensitive to both drugs in our *in vivo* system, with GI50 levels at or below 0.1 µM for most lines, in contrast to 0.1-5 µM *in vitro* (Figure 3A and B). This suggests quite distinct drug sensitivities between the model systems and/or additional drug sensitivity of the tumours being induced by embryonic zebrafish host interactions.

### MRTKIs regorafenib and infigratinib block neo-vascularisation in RMS tumour xenografts

Importantly, for the MRTKIs regorafenib and infigratinib, effect on tumour growth is only one element of their potential mechanism of action, as both drugs have the potential to block growth factor signalling on stromal cells, particularly the vasculature. We, therefore, next assessed the effect of these drugs on the ability of RMS tumours to induce neo-vascularisation by analysing neo-vessel lengths and neo-vessel induction efficiency alongside SIV length and – as control – ISV length, in xenografts treated with 0.1 µM regorafenib, infigratinib, or vehicle (Figure 4, Supplementary figure 7 and 8). This analysis revealed that both drugs had a significant effect on tumour neo-vessel induction (Figure 4A-C, Supplementary figure 7A), with the FP Rh41 line associated vessels particularly strongly affected by regorafenib (complete ablation) and the FN RD line vessels most potently inhibited by infigratinib (2.8% of vehicle treated). The strong effect of regorafenib on Rh41 neo-vessels cannot solely be explained by the fact that Rh41 have the shortest neo-vessels in vehicle treated conditions (Figure 2B), as infigratinib – although effective in inhibiting neo-vessel length across lines, does not have the same striking effect on Rh41. By assessing the proportion of xenografts with neo-vessels present under MRTKI treatment (Figure 4B) and the respective length of these neo-vessels relative to vehicle treatment (Figure 4C), we can dissect the impact on neo-vessel induction and elongation efficiency. This revealed, that regorafenib is far more potent in preventing vessel induction in FP-RMS lines than the FN-RMS RD and JR1 lines (% with neo-vessels relative to DMSO treated: 14.6% vs. 40.9%, respectively). Of note, neo-vascularisation was strongly inhibited (6.4% of vehicle treated) in the FN-RMS line SMS-CTR (Figure 4B). The effect of regorafenib on the neo-vessel length of the other two FN-RMS lines (RD and JR1) seems in large part rendered through inhibition of neo-vessel elongation (19.8% and 15.4% of vehicle treated, respectively) (Figure 4C). The effect of infigratinib, conversely, was primarily in preventing neo-vessel induction, with less impact on neo-vessel elongation (Figure 4B vs. C). Whilst infigratinib was, however, overall less effective in inhibiting neo-vascularisation than regorafenib by both metrics, it importantly inhibited the neovascularisation in FN cell lines to a similar extent than in FP cell lines. Both drugs significantly inhibited the ability of RMS tumours to induce increased SIV proximal bed vascularisation in the majority of lines, with the effect of regorafenib being greater, as SIV lengths was essentially suppressed back to baseline (ie vehicle only injected levels) (Supplementary figure 7B and 8A). Of note, the tumour-distal ISV bed vessel lengths were unaffected by regorafenib or infigratinib, suggesting the effects seen by drug treatment on tumour-proximal vessel metrics were again due to modulation of the short-range tumour-vessel signalling (Supplementary figure 8B).

**Figure 4:**
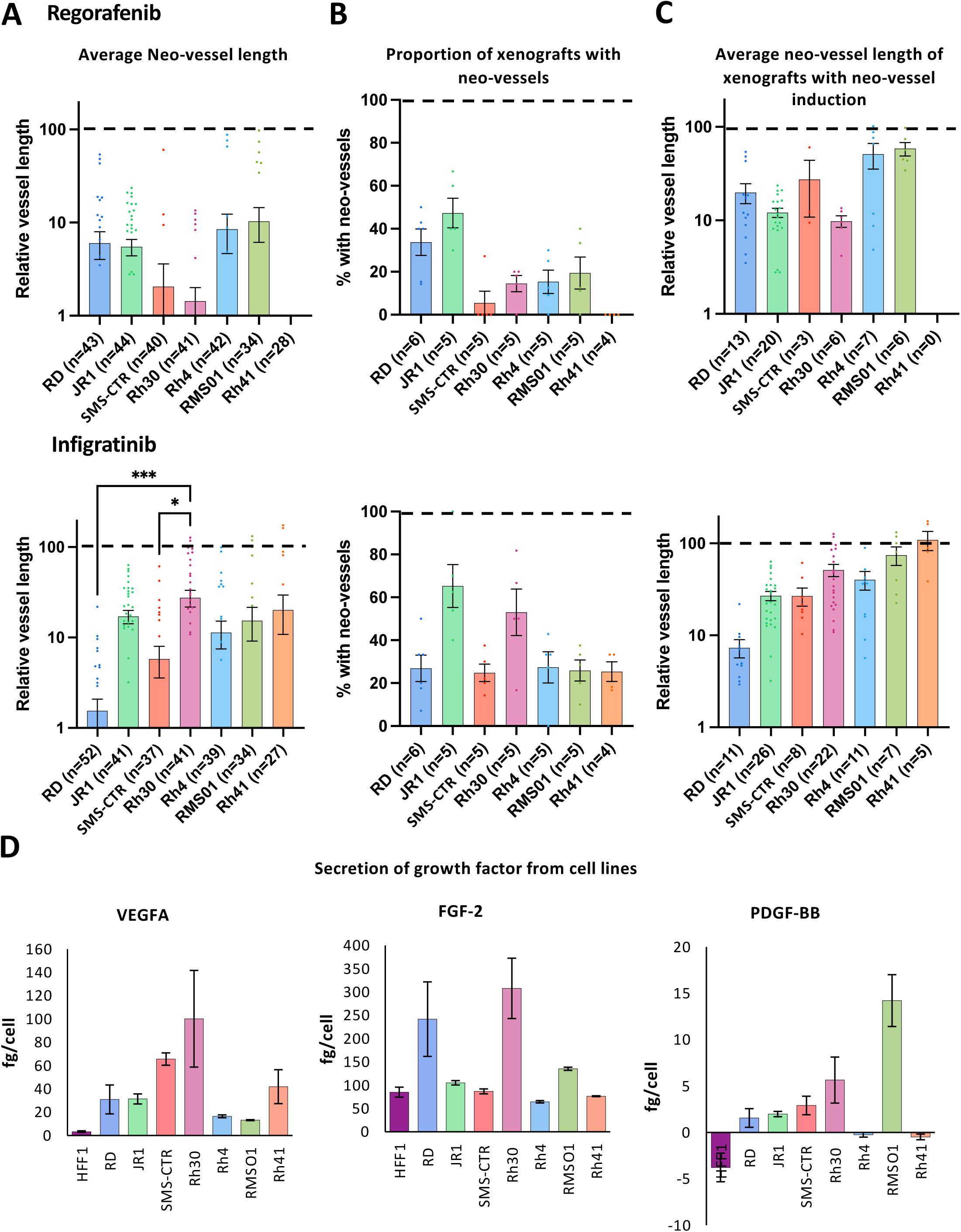
Regorafenib and infigratinib treatment inhibits RMS xenograft tumour induced vascularisation. Xenografts generated from the 7 RMS cell lines were treated with 0.1 µM regorafenib or infigratinib for 66 hours after which vessel development was assessed by FIJI image analysis. **A**, Bar graph of the effect of each treatment on tumour induced neo-vessel length overall, relative to the average vehicle treated vessel length for each cell line (dotted line 100%). **B**, Bar graph of the proportion of xenografts with tumour induced neo-vessels, upon drug treatment. **C**, Bar graphs of average length of tumour induced neo-vessel, of those where neo-vessels were induced, relative to the average vehicle treated vessel length for each cell line. * P<0.05, ** P<0.01, *** P<0.001, **** P<0.0001, Ordinary one-way ANOVA with Tukey’s multiple comparison’s test, Data are presented as mean values +/− SEM. **D**, Bar graphs showing the amount of secreted VEGFA, FGF-2 and PDGF-BB secreted in 24hrs *in vitro* by each cell line, assessed by ELISA of conditioned media. n = number of xenografts (except for B, which is number of experiments).

Overall, analysis of MRTKI drug responses revealed significant heterogeneity between RMS tumours in terms of the level of drug-induced vessel disruption. As both, regorafenib and infigratinib target pro-angiogenic growth factor receptors (VEGFRs/PDGFR and FGFRs), we hypothesised that this heterogeneity could be due to variations in the production of growth factor substrates by the tumour cells. To investigate this, we performed ELISA analysis for the primary pro-angiogenic substrates for each of these receptors (VEGFA, PDGF-BB and FGF-2), in tumour-conditioned media, with growth factor production from HFF1 cells also assessed by way of non-tumourigenic reference (Figure 4D). Strikingly, this analysis revealed an inverse correlation between VEGFA secretion and neo-vessel length upon regorafenib treatment, between RMS lines (R=-0.76, P=0.05), indicating VEGFA secretion as a potential marker of vessel response to regorafenib (Figure 4D). The link between FGF-2 production and vessel inhibition by infigratinib was less clear; however, the relatively high production of FGF-2 in RD and Rh30 and the potent inhibition of the neo-vessel and SIV bed induction in these lines by infigratinib, could suggest a role for this drug in FGF-2 induced angiogenic signalling blockade (Figure 4D).

To further explore the interrelatedness of our functional analysis data, we performed Pearson correlation analysis between fusion status, angiogenic growth factor secretion, tumour and vessel growth metrics and response to MRTKis regorafenib and infigratinib for all cell lines (Supplementary figure 9). This analysis confirmed the significant correlation between FP status and larger tumour size in our xenografts (R=0.73, P=0.04), as well as the correlation of VEGFA secretion and neo-vessel induction sensitivity upon regorafenib treatment. It, additionally, revealed that the pattern of effect of both, regorafenib and infigratinib, on tumour size is significantly correlated (R=0.83, P=0.01) and that tumours secreting higher PDGF-BB drive a greater level of SIV elongation (R=0.91, P=0.004) (Supplementary figure 9).

### Human growth factor injection induces neo-vessels which phenocopy tumour vessel drug responses

Given the striking effects of yolk-injected tumour cells on neo-vessel formation and extended SIV formation which can be modulated by the MRTKIs, we hypothesised that the introduction of RMS cells into the yolk of embryonic zebrafish induces non-physiological blood vessel sprouting from the SIV bed and SIV bed extension through the production of pro-angiogenic growth factors. We have demonstrated a potential role for VEGFA, FGF-2 and PDGF-BB in mediating these dynamics. In order to validate that human growth factors are capable of inducing embryonic zebrafish neo-vessel formation we injected recombinant human VEGFA, FGF-2 or PDGF-BB into the yolk of zebrafish embryos at 50 hpf (mirroring the time when tumour cells would be injected) and quantified neo-vessel and SIV lengths after 70hrs (Supplementary figure 10). This analysis identified that a growth factor dose of 0.5 and 1.5ng for VEGFA and FGF-2, respectively, was sufficient to induce neo-vessels in >80% of embryos and at an average length of 141 and 146 µm, similar to those induced by the RMS tumours (36-166 µm) (Supplementary figure 10A-C compared to Figure 2B&C). Lower doses of VEGFA and FGF-2 (0.17 and 0.5ng, respectively) induced neo-vessels in only 39 and 32% of embryos, respectively (Supplementary figure 10A). These results are consistent with previous studies demonstrating the ability of exogenous VEGF to induce neo-vascularisation in zebrafish (41–43). PDGF-BB on the other hand induced only sporadic neo-vessels at a dose of 1.5ng (Supplementary figure 10A) and no neo-vessels at a dose of 0.5ng (Supplementary figure 10A). As our previous analysis of PDGF-BB production by RMS cells found it to be in the fg/cell range (Figure 4D), we decided any further dose escalation of PDGF-BB would not be physiologically relevant. SIV lengths were also significantly increased by all growth factors to a similar degree to that seen in the RMS tumour xenografts (% SIV length increase over vehicle injected; 39% VEGFA, 26% FGF-2, 35% PDGF-BB) (Supplementary figure 10D compared to Figure 2C). The observation that PDGF-BB injection specifically induces SIV elongation, but not neo-vascularisation is in agreement with our previous finding that the level of PDGF-BB secretion by RMS tumour lines is significantly correlated with SIV elongation in our xenografts (Supplementary figure 9). Importantly upon treatment with 0.1 µM regorafenib or infigratinib, tumour growth factor induced zebrafish angiogenesis was blocked, further supporting the case for the involvement of these growth factors in mediating drug-induced neo-vessel response. Intriguingly, both, regorafenib and infigratinib, were capable of blocking VEGFA, PDGF-BB and FGF-2 induced angiogenesis, albeit regorafenib showed greater selectivity for VEGFA and PDGF-BB induced angiogenesis and infigratinib for FGF-2 induced angiogenesis, suggesting a level of cross-talk between the growth factors and their respective pathways in the induction of angiogenesis (Supplementary figure 10).

### Patient-derived FP RMS cells grow and vascularise in the embryonic zebrafish and shows response to MRTKIs

We have demonstrated the capacity of the here developed embryonic zebrafish xenograft model to investigate and segregate the effects of MRTKIs on RMS tumours and their accompanying vasculature when they were generated from cell lines. The behaviour of cell lines, however, often diverges significantly from clinical tumours due to epigenetic drift and clonal selection of tumour cells best suited to non-physiological culture conditions (reviewed in(44)). For this model to optimally contribute to our understanding of clinical RMS and its treatment responses, it is important that it can model the behaviour of tumour cells with a close to clinical identity. We, therefore, assessed whether patient-derived FP-RMS cells IC-pPDX-104 which are well characterised and had previously been assessed with regorafenib treatment *in vitro* and *in vivo* (45), were able to grow and induce neo-vessel formation in our model system (Figure 5A). This analysis revealed that IC-pPDX-104 cells form on average significantly larger tumours in the xenograft model than the cell lines (average tumour area; 16951 µm^2^ FN-RMS, 21989 µm^2^ FP-RMS, 26074 µm^2^ IC-pPDX-104; p<0.0001 and p<0.0086 versus FN-RMS and FP-RMS respectively) (Figure 5A, Supplementary figure 4, compared to Figure 1D and 5A), induce neo-vessels with an efficiency of 93% and of an average length of 82.7 µm, both comparable to the RMS cell lines (94.5% and 127.9 µm FN-RMS, 81.1% and 71.2 µm FP-RMS) (Supplementary figure 7A, compared to Figure 2B and 5A (2^nd^ and 3^rd^ panel) and significantly enhance the length of the proximal SIV bed by on average 27.2% compared to vehicle only injected embryos, again in line with the 24.5% and 29.1% increase induced by FN– and FP-RMS cell lines, respectively (Supplementary figure 7B, compared to Figure 2C and 5A (4^th^ panel)). Like all the other RMS tumour samples, IC-PDX-104 did not significantly affect the ISV length in the xenografts compared to vehicle injected controls (Data not shown).

**Figure 5:**
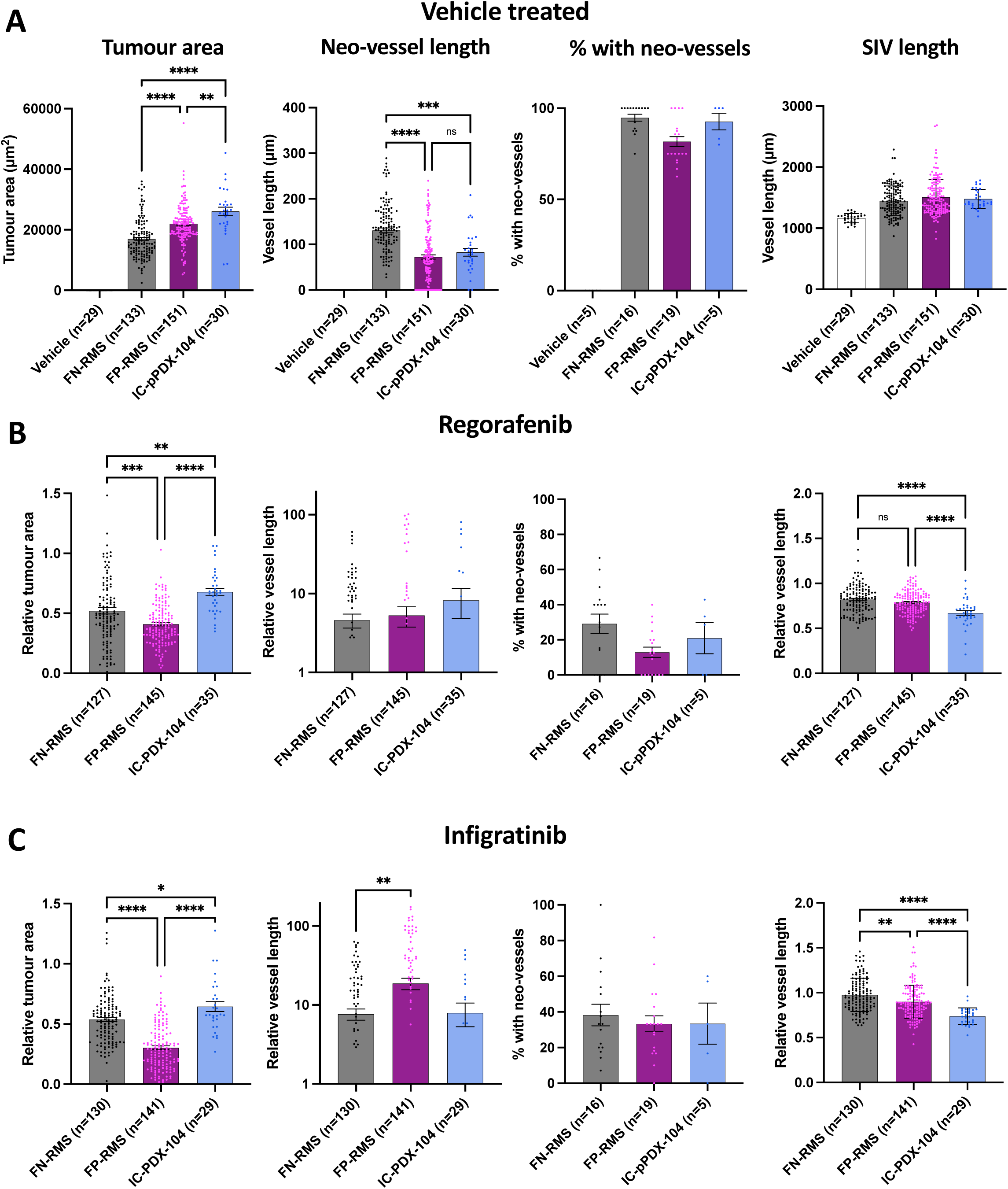
Patient derived RMS cells derived from an in mouse established xenograft also form vascularized tumours in the zebrafish xenograft setting, which are responsive to MRTKI inhibition. **A**, Bar graphs showing tumour area, neo-vessel length, neo-vessel frequency and SIV length for vehicle only injected as well as all FN, all FP and the FP IC-pPDX-104 injected xenografts. **B**, Bar graphs showing the effect of 0.1 µM regorafenib treatment on tumour area, neo-vessel length, neo-vessel frequency and SIV length for all FN,, all FP and FP IC-pPDX-104 injected xenografts. **C**, Bar graphs showing the effect of 0.1 µM infigratinib treatment on tumour area, neo-vessel length, neo-vessel frequency and SIV length for all FN, all FP and FP IC-pPDX-104 injected xenografts. * P<0.05, ** P<0.01, *** P<0.001, **** P<0.0001, Ordinary one-way ANOVA with Tukey’s multiple comparison’s test, Data are presented as mean values +/− SEM. n = number of xenografts (except for “% with neo-vessels”, which is number of experiments).

Furthermore, similarly to cell lines, inhibition with the MRTKIs regorafenib and infigratinib significantly reduced tumour area, as well as neo-vessel and SIV induction and elongation in the embryonic zebrafish model system (Supplementary figure 4 and 7A&B, compared to Figure 5B, C). The effect on tumour size was clear, but less pronounced than in our cell line models (relative tumour area; 0.68 and 0.64 IC-pPDX-104, 0.52 and 0.52 FN-RMS and 0.41 and 0.29 FP-RMS, regorafenib and infigratinib respectively). This aligns with a lesser *in vitro* responsiveness of IC-pPDX104 to regorafenib in particular (average GI50; 9.1 µM and 1.8 µM IC-pPDX-104, 3.5 µM and 2.6 µM FN-RMS, 2.0uM and 1.0uM FP-RMS, regorafenib and infigratinib respectively) (Supplementary figure 6) and with a lack of tumour growth reduction for single agent regorafenib at 15 mg/kg daily orally of subcutaneous xenografts in mice (45) whereas RMS01 subcutaneous xenografts showed a slowing of tumour growth with 10mg/kg, as well as 30 mg/kg daily regorafenib treatment in another study (16). Overall, these findings suggest that our embryonic zebrafish xenograft model system can be used to investigate treatment response dynamics in patient-derived tumours, highlighting the potential for clinical translation of this model system.

## Discussion

RMS is a very aggressive soft tissue sarcoma of childhood and young adulthood and – despite international efforts of expert consortia – we still struggle to a) readily identify promising new treatments for introduction into clinical trials and b) identify promising biomarkers to predict response to therapy (conventional or novel). To advance in these areas, it is essential to optimise our preclinical assessment tools and – if considered required – develop new preclinical models as undertaken here in our study with the development of the embryonic zebrafish xenograft model.

Up to now, the majority of pre-clinical modelling in RMS has utilised either *in vitro* 2D and 3D cultures or assessed RMS tumour growth *in vivo* mostly using RMS cell lines or PDX implanted into mice subcutaneously or intramuscularly (46). These models are poorly adapted for investigating tumour-stromal interactions – including with the vasculature – and the effect of therapeutics on these. Recently, the juvenile and adult zebrafish has been added to the armamentarium of RMS research to study both implanted and genetically or chemically induced RMS (46–52), but with neovascularisation and existing vessel bed expansion not formally investigated and still challenging to accomplish due to the limited translucency of the zebrafish, even when using pigment deficient transgenic forms such as the *roy*^-/-^/*nacre*^-/-^ (*Casper*) zebrafish or its immunodeficient variant *Casper prkdc^-/-^*, *il2rga^-/-^* (to allow xenografting of the adult zebrafish) (53, 54). The latter publication developed several cancer xenografts including RMS xenografts of EGFP^+^ cell lines and EGFP^+^ patient-derived cells. The RMS xenografts were either injected into the peritoneal cavity for tumour growth and treatment studies, or injected peri-ocularly allowing single cell imaging analysis of different tumour populations through the translucent skin (54). Whilst this can undoubtably provide new insights into the behaviour of RMS, it so far cannot optimally assess tumour-vessel interactions.

Over the more recent years, embryonic zebrafish models gained traction in cancer research, due to various reasons – a few summarized here; embryonic zebrafish are translucent, easy to genetically manipulate – such as *flk1::GFP* transgenes leading to green colour of the endothelial cells – and provide the possibility to create cancer xenografts and follow their growth and physiological and pathological angiogenesis with advanced imaging techniques in high throughput (30, 32, 55). Several different cancer cell lines including osteosarcoma and Ewing sarcoma cell lines have been successfully engrafted so far and limited studies of lung, colon and breast cancer cell lines had explored the impact of xenografts and treatment with anti-angiogenics on vessel formation in the embryonic zebrafish (56, 57). However, systematic assessment of clinically relevant drugs affecting tumour stroma interaction with focus on angiogenesis has not been published for any tumour type and especially RMS cell lines or xenografts have not yet been studied apart from in a recent limited study which injected RMS cell lines RD, Rh30 and Rh5 at 3hpf blastula stage – without anatomical structures yet discernible; only RD engrafted and vessel induction was not assessed (58).

Our manuscript details the development and testing of a new embryonic zebrafish xenograft model system for assessment of RMS tumour development, tumour-vessel interactions and drug responses on tumour growth and vessel formation. We establish injection at 50 hpf and injection in the yolk as most successful conditions for further analysis. We demonstrate that tumours from a range of RMS cell lines and a patient-derived sample are viable, grow and induce neo-vessels and expand the existing SIV in the model system. This is noteworthy as the tumour cells are grown at 34°C in the model system, a compromise between the 37°C human and 28.5°C zebrafish optimum temperatures to maximise viability in both components of the model system. We demonstrate the ability of this model to dissect distinct tumour and associated vessel response profiles to clinically relevant MRTKIs, regorafenib and infigratinib, addressing an unmet need in the field for models to allow better understanding of the primary mechanism of action of multi-targeted therapeutics.

We, additionally, demonstrate a strong correlation between the cellular production of the pro-angiogenic growth factor, VEGFA and tumour neo-vascularisation to regorafenib. VEGF expression has previously been found to be associated with poor prognosis in both, FN and FP RMS(59), with both VEGF and PDGF found to be the main drivers of angiogenesis in RMS, alongside hypoxia (59, 60). In addition to the production of pro-angiogenic growth factors by RMS cell lines, we also have examined the effect of these factors on neo-vessel induction in the tumour-free embryonic zebrafish. Intriguingly, the calculated growth factor production from the RMS cell lines is at maximum 100, 308 and 14 fg/cell for VEGFA, FGF-2 and PDGF-BB, respectively. Therefore, in the zebrafish xenograft model system assuming, based on quantifications detailed in Figure 1, an average of 921 cells present over the 70 hours of the experiment growth factor production would be 92, 284 and 13 pg, which is well below the injected growth factor amount (500-1500pg; detailed in Supplementary figure 10) to achieve the same level of neo-vascularisation. One possible explanation for this would be the synergistic relationship between pro-angiogenic paracrine signalling, with RMS cells producing a variety of growth factors supporting vessel induction, not all of which have been profiled in this study. Therefore, far less of each growth factor may be needed to achieve neo-vascularisation than a single injected growth factor. The reduced temperature of 34°C experienced by the RMS cell in the xenograft may also impact growth factor production and contribute to this disparity. Xenografted tumour cells of a glioma have also previously been shown to induce the ectopic production of zebrafish pro-angiogenic growth factors including VEGFA and its receptor VEGFR2, accompanied by increased SIV branching (43). If a similar process is occurring in RMS, this could also contribute to the vascularisation of these tumours. Additionally, PDGF-BB has been shown to modulate angiogenesis by stimulating VEGF and FGF production in endothelial cells (61, 62). The vascularisation dynamics seen in our model is likely to be the result of significant crosstalk between multiple pro-angiogenic growth factors representing a possible explanation why regorafenib and infigratinib were able to inhibit both VEGFA and FGF-2 recombinant growth factor induced neo-vascularisation, despite having limited direct inhibitory capacity to their receptors (FGF-2 in the case of regorafenib and VEGFA in the case of infigratinib) (Supplementary figure 10). These data support further the clinical approach to study MRTKI such as regorafenib in RMS and can add an explanation to the negative result of the Phase II randomised, clinical trial in metastatic RMS in frontline comparing standard of care chemotherapy with standard of care chemotherapy with the addition of Bevacizumab, a humanized monocloncal antibody against VEGF-A(63). Our findings suggesting that the level of secretion of VEGFA mediates drug response to regorafenib could be the first step of identifying a biomarker for response to regorafenib and it would be interesting to assess particularly VEGF-A, but also FGF-2 and PDGF-BB in tumour samples in the clinical context such as our FaR-RMS platform trial in the relapse setting where two therapeutic regimens – vincristine, irinotecan and temozolomide versus the experimental arm of vincristine, irinotecan and regorafenib are investigated in randomized fashion (2).

The drug response data detailed in this study overall suggests that regorafenib and infigratinib have potentially quite distinct mechanisms of action in our model of RMS tumours, with heterogeneity in infigratinib drug response primarily associated with distinct tumour cell sensitivities, as suggested by the similar pattern of response seen in the tumour cell only *in vitro* model system. This suggests that blockade of autocrine signalling is critical for tumour response to infigratinib. Heterogeneity in regorafenib response on the other hand may also be a product of differential sensitivities to paracrine tumour-vessel signalling inhibition, as demonstrated by the striking correlation of regorafenib induced tumour neo-vessel inhibition and production of VEGFA by the respective cell lines. The direct tumour cell targeting (autocrine-blockade) effect of infigratinib, particularly in FP RMS has previously been demonstrated by Milton et al., 2022 (19), who highlighted the importance of FGF7-FGFR2 autocrine signalling in treatment response. Our study corroborates this, but importantly also highlights the potential for infigratinib to block paracrine tumour vessel production, as demonstrated by the significant inhibition of neo-vessel induction in all RMS cell lines tested and the IC-pPDX-104, but particularly in the FN lines RD and SMS-CTR. This suggests the intriguing possibility that infigratinib could have distinct primary anti-tumour mechanisms in different RMS populations, directly inhibiting autocrine signalling induced tumour growth in FP RMS and acting as paracrine-signal blocking anti-angiogenic agent in some FN lines. This demonstrates a previously unappreciated potential for infigratinib to act as an anti-angiogenic agent in RMS, particularly in the FN sub-type where direct anti-tumour activity had previously been shown to be disappointing (19) and could suggest benefit of infigratinib for FN RMS. These findings further highlight the limitations of tumour cell only *in vitro* drug response assays and point towards the importance of modelling tumour-stromal drug responses.

In addition to RMS cell lines, we also tested the ability of patient-derived RMS tumour cells of one patient to grow and vascularise in embryonic zebrafish xenografts, demonstrating that they produce viable and reproducible tumours with similar growth and vascularisation characteristics to RMS cell line tumours and similar responses to regorafenib and infigratinib. This demonstrates the translatability of these findings to preclinical testing of patient-derived tumour cells.

As a further aspect of this study, we identify the yolk region as a more effective site than the PVS for RMS tumour cell seeding in zebrafish embryos. This is surprising as the PVS is becoming increasingly favoured as the preferable site for tumour cell inoculation, due to improvements in tumour growth over yolk inoculation (55, 64). We find the opposite to be true in RMS. It should be noted that most previously published zebrafish xenograft models and those where the PVS is favoured have been modelling carcinomas which are quite distinct in their cell of origin, signalling and behaviour to sarcomas, such as RMS (65, 66). It further is in line with the finding that the embryonic zebrafish yolk has been characterized, as a hypoxic, avascular space, which as such may provide an ideal environment for RMS which is hypoxia prone (67).

The ability to profile the initial steps of vessel induction and the effect of therapeutics on this, sets it apart from mouse-based *in vivo* modelling, that rather looks at the later stage effects of the drugs on tumour growth and vascularisation. Both, early and late stage effects of therapeutics, are important to model. The focus of the zebrafish xenograft model on a short time window of the initial steps of tumour establishment and vascularisation from a small number [250] of cells, suggests the system could potentially be used to investigate the efficacy of therapeutics against residual RMS cells after intensive chemotherapy, at a stage where maintenance chemotherapy is currently given in the clinic in high-risk patients. This would serve investigations to improve maintenance therapy and to study prevention of relapse. Preliminary experiments suggest that as few as 32 RMS cells can proliferate and induce angiogenesis in the embryonic zebrafish yolk (data not shown). The activity of regorafenib and infigratinib to block RMS tumour paracrine signalling induced neo-vascularisation, demonstrated in our study, could support the use of these drugs in the tumour maintenance setting.

Further systematic analysis of patient samples is required in our model; it could be undertaken as part of a wider testing alongside established models like 2D and 3D culture and *in vivo* PDX xenograft modelling in mice, this would allow to directly compare the different preclinical model systems. Parallel analysis in different models will likely provide a more comprehensive picture of the effect of therapeutics. Given the varying outcome measures between different models – e.g. GI50 in 2D culture, % growth reduction and % vessel lengths reduction in embryonic zebrafish, tumour volume in mouse xenografts – it will be important to define thresholds for each model system which will be indicating positive responses to therapeutics; to accomplish this, it will be necessary to combine preclinical testing of patients’ tumour cells with the parallel clinical assessment of drugs in the same patients (independent of the outcome of the preclinical testing) as part of a clinical trial with clinical response data available to researchers in an anonymized manner; i.e., a co-clinical trial setting (Figure 6) which could be initiated as part of the currently ongoing relapse study in FaR-RMS assessing regorafenib (2). As we still commonly fail to predict responses to therapy in the clinic based on preclinical modelling this approach would be desirable and is necessary to norm and validate our preclinical tools, ie to allow to determine the individual, specific value of each model system in predicting responses in the clinic and in our case, to specifically define the preclinical place for the here developed embryonic zebrafish model (Figure 6).

**Figure 6:**
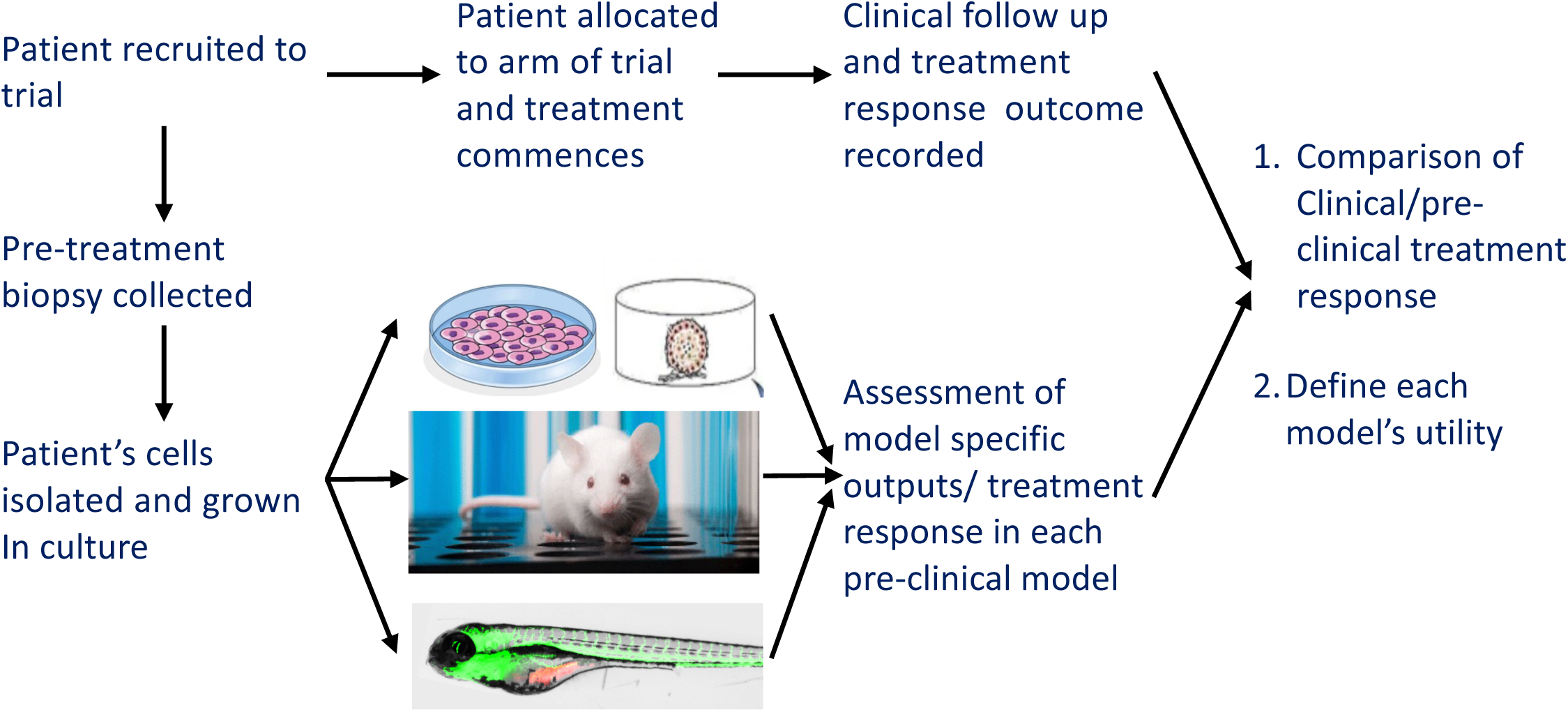
Schematic illustrating the proposed testing regime for pre-clinical modelling options in a co-clinical trial. The various aims and benefits are outlined in the figure.

## Data availability statement

The study did not generate new unique reagents. The data generated in this study are available within the article and Supplementary Data.

## Author contributions

JWW: Project conceptualisation and coordination, Methodology, Data creation and curation, Data analysis and interpretation, Funding acquisition, Writing – writing original draft and review/editing. ELG: Methodology, Data creation and curation, Writing – original draft and review/editing; R.M.: Zebrafish maintenance, Data interpretation, Writing – original draft and review/editing. JRM: supported tissue culture maintenance of the RMS cell lines, Data interpretation, Funding acquisition, Writing – original draft and review/ editing. ADB: supported imaging of the xenograft model and cell culture systems, Writing – original draft and review/editing. FM: Data interpretation, Funding acquisition, Writing – original draft and review/editing. SAG: Project conceptualisation and coordination, Data interpretation, Funding acquisition, Supervision, Writing – original draft and review/editing.

## Supporting information

Wragg et al., Supplementary figures

## Acknowledgements

We are extremely grateful to Sara and David Wakeling, the founders of Alice’s arc for their support and their ongoing enthusiasm for this work. The RMS cell lines were kindly provided by Prof. Janet Shipley, ICR, Sutton, UK. We are grateful to Prof. Olivier Delattre, for the generation of the patient-derived cell line IC-pPDX-104, and to Prof. Beat Schäfer and Dr. Marco Wachtel for sharing it with us in agreement with Prof. Delattre. We also would like to thank Prof. Beat Schäfer and Dr. Marco Wachtel, Universitätsspital Zürich, Switzerland for helpful discussions and the University of Birmingham Biomedical Services Unit for their support with the supply and maintenance of zebrafish lines for this project. Part of the methodology of this work has been developed in the thesis of ELG (Master of Research in Cancer Sciences at the University of Birmingham).

