## Supplementary material for "A dual readout embryonic zebrafish xenograft model of rhabdomyosarcoma to assess clinically relevant multi-receptor tyrosine kinase inhibitors": Wragg et al., Supplementary figures

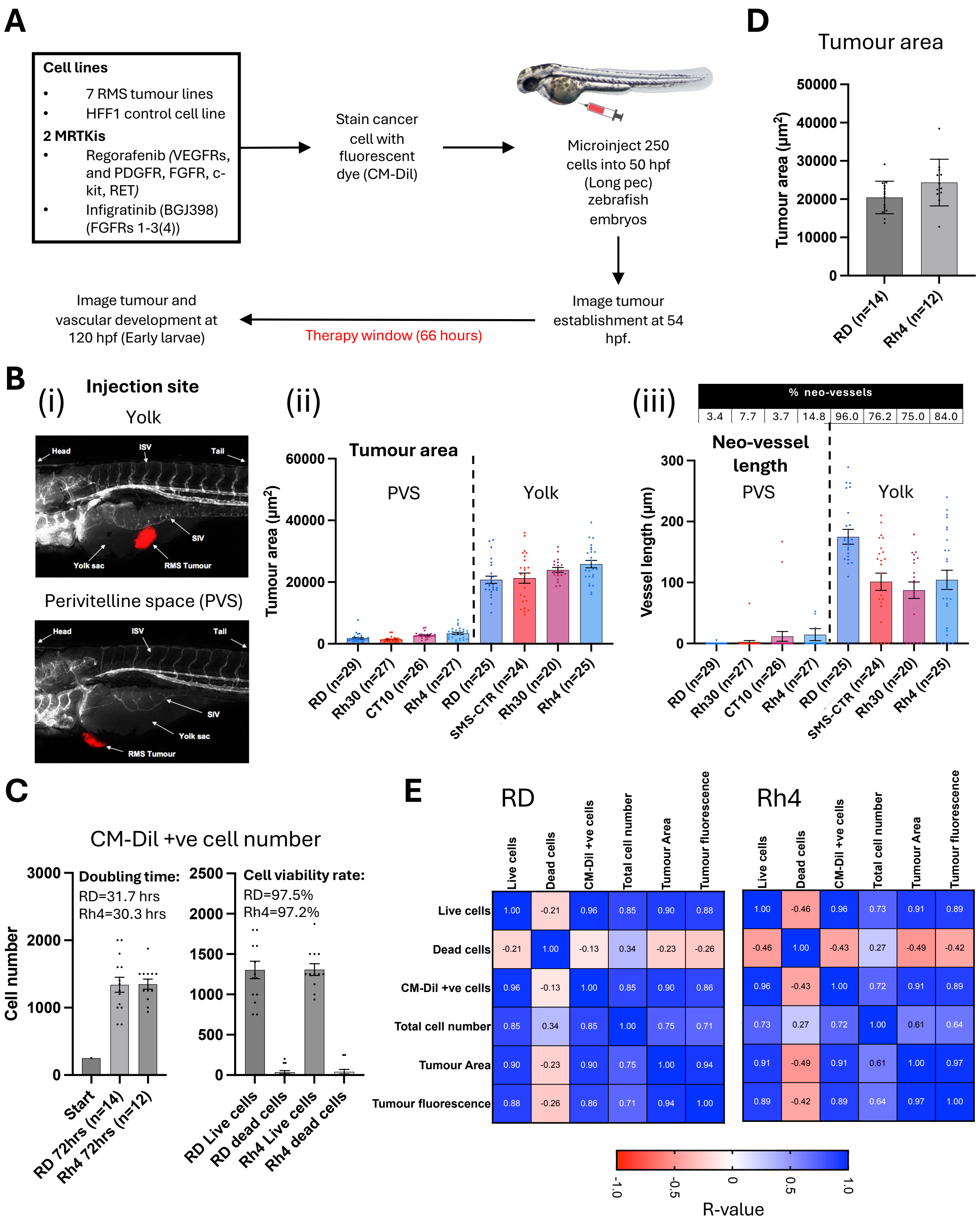

**Supplementary figure 1: RMS tumour cells are viable and proliferate in larval zebrafish hosts.**

**Supplementary figure 1: RMS tumour cells are viable and proliferate in larval zebrafish hosts**. **A,** Schematic of the xenograft model setup workflow with hours post fertilisation (hpf) and developmental stages described. **B**, Comparison of xenograft injection sites for RMS cells. **(i)** images of yolk vs. PVS injected tumours, imaged at 120 hpf [Early larvae] (end of experiment). **(ii)** bar graph of tumour area of PVS and yolk injected tumours. **(iii)** bar graph of neo-vessel length including at the top of the graph a table of neo-vessel frequency of PVS and yolk injected tumours, measured as described in Figure 3A. Data are from 4 RMS lines and 3 experiments where the two injection sites were directly compared. **C**, Bar chart showing total CM-Dil positive (left) and CM-Dil positive live/dead (right) cell number counts from excised RD and Rh4 tumours, measured at the end of the xenograft experiment (70 hpi) and compared to the initial injected bolus of 250 cells. Doubling time and cell viability rate are shown. **D**, Tumour area was calculated from fluorescent images of the xenografts measured for cell counting as described in Figure 2A. **E**, Pearson correlation matrix between live, dead, CM-Dil positive and total cell numbers calculated by cell counting with tumour area and fluorescence calculated by image analysis from RD and Rh4 tumours. R-values are shown. Data are presented as mean values +/- SEM. n = number of xenografts.

**A**

| | RD (n=51) | JR1 (n=42) | SMS-CTR (n=40) | Rh30 (n=38) | Rh4 (n=40) | RMS01 (n=39) | Rh41 (n=34) |
|---|---|---|---|---|---|---|---|
| Neo-vessels induced | 50 | 42 | 34 | 32 | 36 | 31 | 24 |
| No neo-vessels | 1 | 0 | 6 | 6 | 4 | 8 | 10 |
| Average neo-vessel length all xenografts (µm) | 165.75 | 122.33 | 95.63 | 85.50 | 110.71 | 52.58 | 35.90 |
| Average neo-vessel length neo-vessel +ve xenografts (µm) | 169.06 | 122.33 | 112.51 | 101.53 | 123.02 | 66.15 | 50.86 |
| % change in SIV length vs. vehicle only injected | 23.49 | 25.27 | 24.82 | 35.55 | 23.59 | 40.74 | 16.89 |

**B**

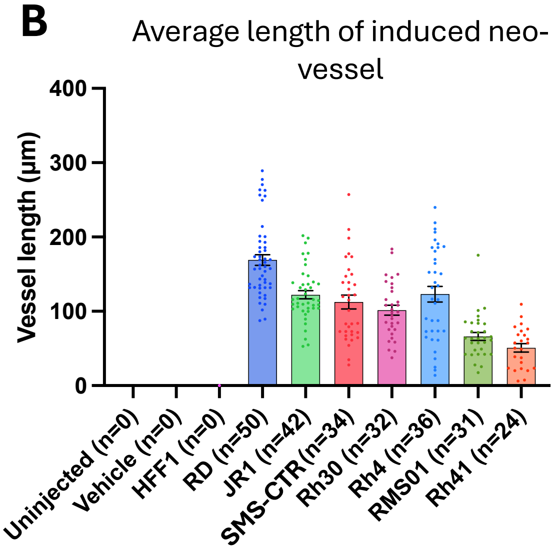

**Supplementary figure 2**: **RMS tumours induce neo-vascularisation from proximal vessel beds. A**, Table of key statistics for neo-vessel induction efficiency, extent of elongation and SIV length change. **B**, bar graph showing the average neo-vessel length of those induced (excluding where no neo-vessels were induced) for xenografts generated from each RMS cell line. Data are presented as mean values +/- SEM. n = number of xenografts.

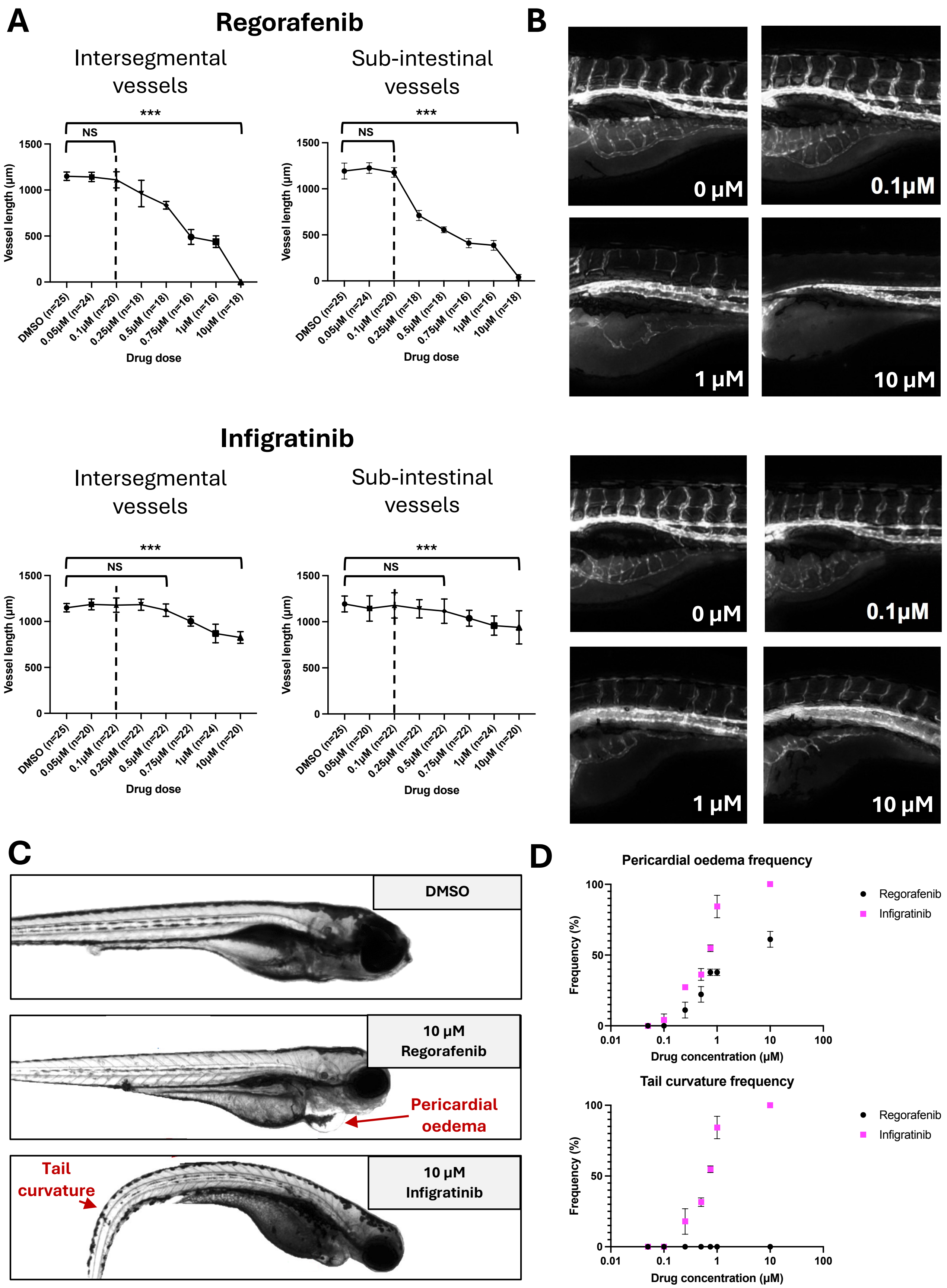

**Supplementary figure 3: Assessing effect of multi-receptor tyrosine kinase inhibitors on tumour free zebrafish embryos**

**Supplementary figure 3: Assessing the effect of MRTKI on tumour free zebrafish embryos.** **A**, 50 hpf *flk1-GFP* zebrafish embryos were exposed to regorafenib or infigratinib drug treatment at a range of concentrations (0.05 μM – 10 μM) and treated for 66 hours prior to assessment of the effect on vessel development by imaging. Line graphs showing ISV and SIV lengths upon treatment. Dashed line shows chosen drug dose of 0.1 μM (*** P<0.001, Ordinary one-way ANOVA with Tukey's multiple comparison's test, data are presented as mean values +/- SD). **B**, Representative images of vessel development at the end point of the drug treatment experiment. **C**, Representative images of drug induced tail curvature and pericardial oedema developmental abnormalities. **D**, Scatter plot of the frequency of tail curvature and pericardial oedema developmental abnormalities upon treatment with regorafenib and infigratinib (data are presented as mean values +/- SEM). n = number of xenografts

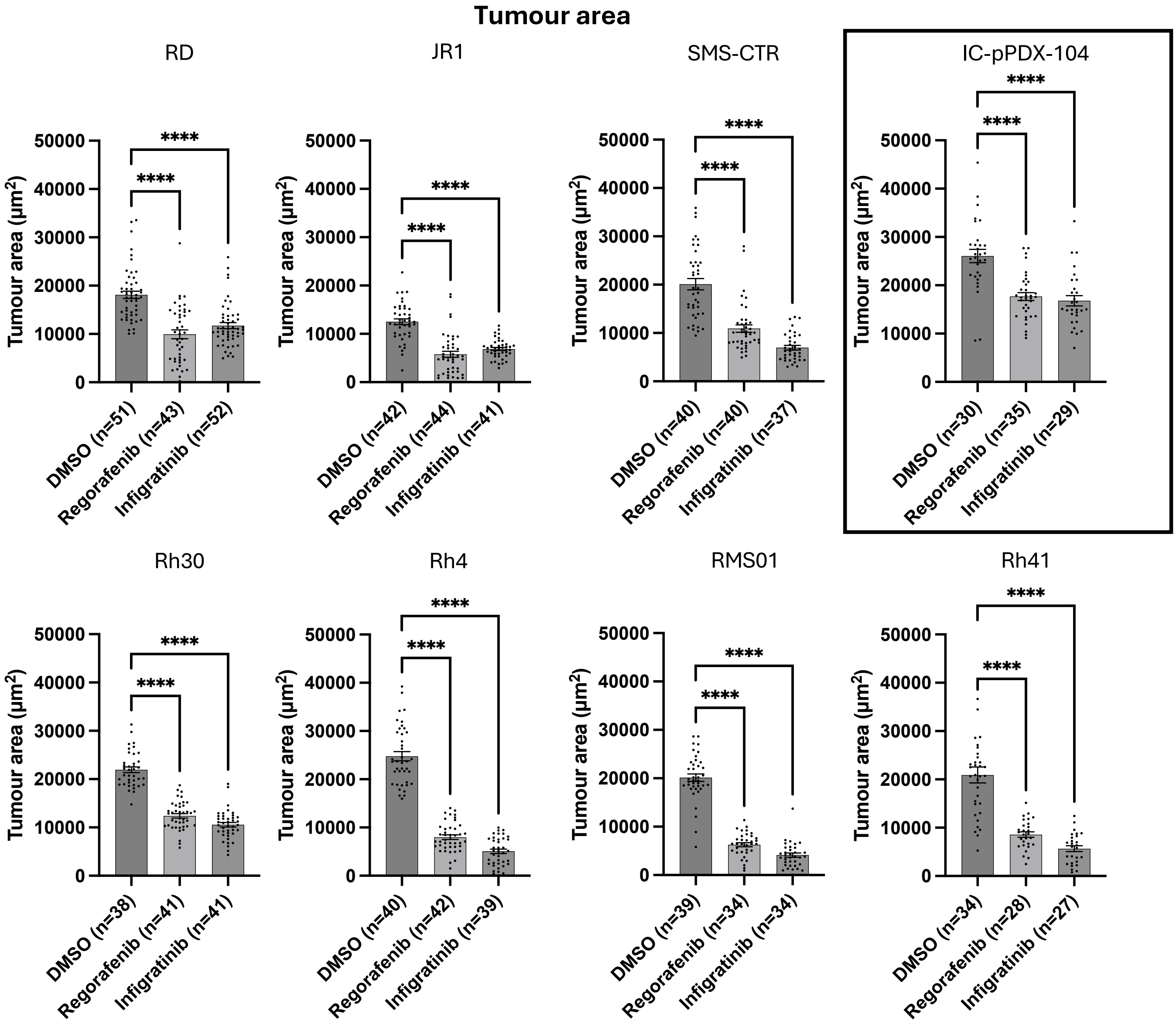

**Supplementary figure 4: Regorafenib and infigratinib treatment inhibits RMS xenograft tumour growth.** Xenografts generated from the 7 RMS cell lines and 1 patient derived culture were treated with 0.1 μM regorafenib or infigratinib for 66 hours. Bar graphs of absolute tumour area values for treated xenografts of each cell line. n = number of xenografts.

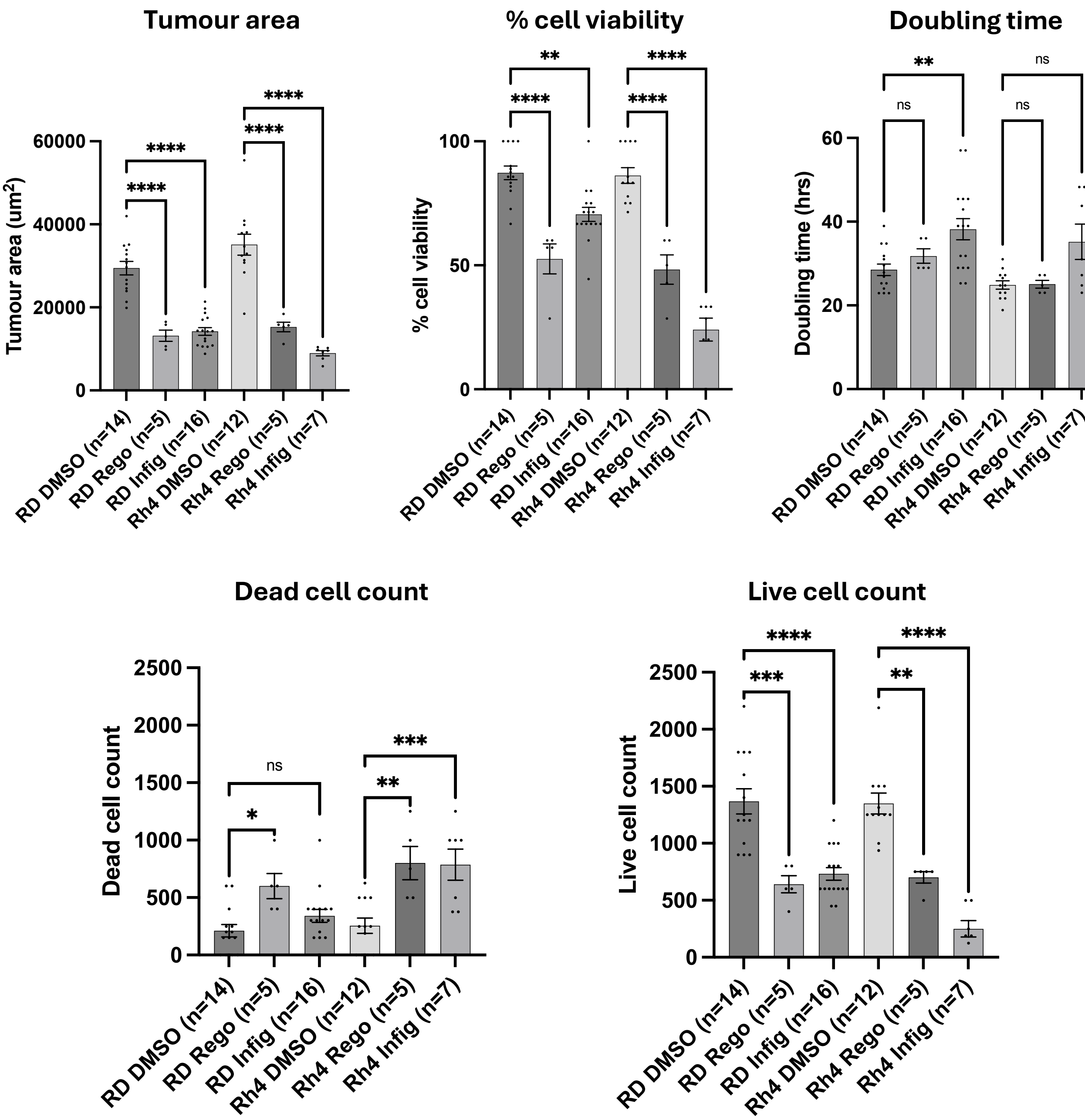

**Supplementary figure 5**: **Regorafenib and infigratinib treatment reduces RMS xenograft viability and proliferation.** Hemocytometer-based cell counting was performed (as described in Figure 1A&B) alongside image-based tumour area measurements for a subset of xenografts generated from RD and Rh4 cell lines. Bar graphs showing drug effect on tumour area (by image analysis) and cell viability, doubling time and live/dead cell counts (from excised tumours). * P<0.05, ** P<0.01, *** P<0.001, **** P<0.0001, Ordinary one-way ANOVA with Tukey's multiple comparison's test, Data are presented as mean values +/- SEM. n = number of xenografts.

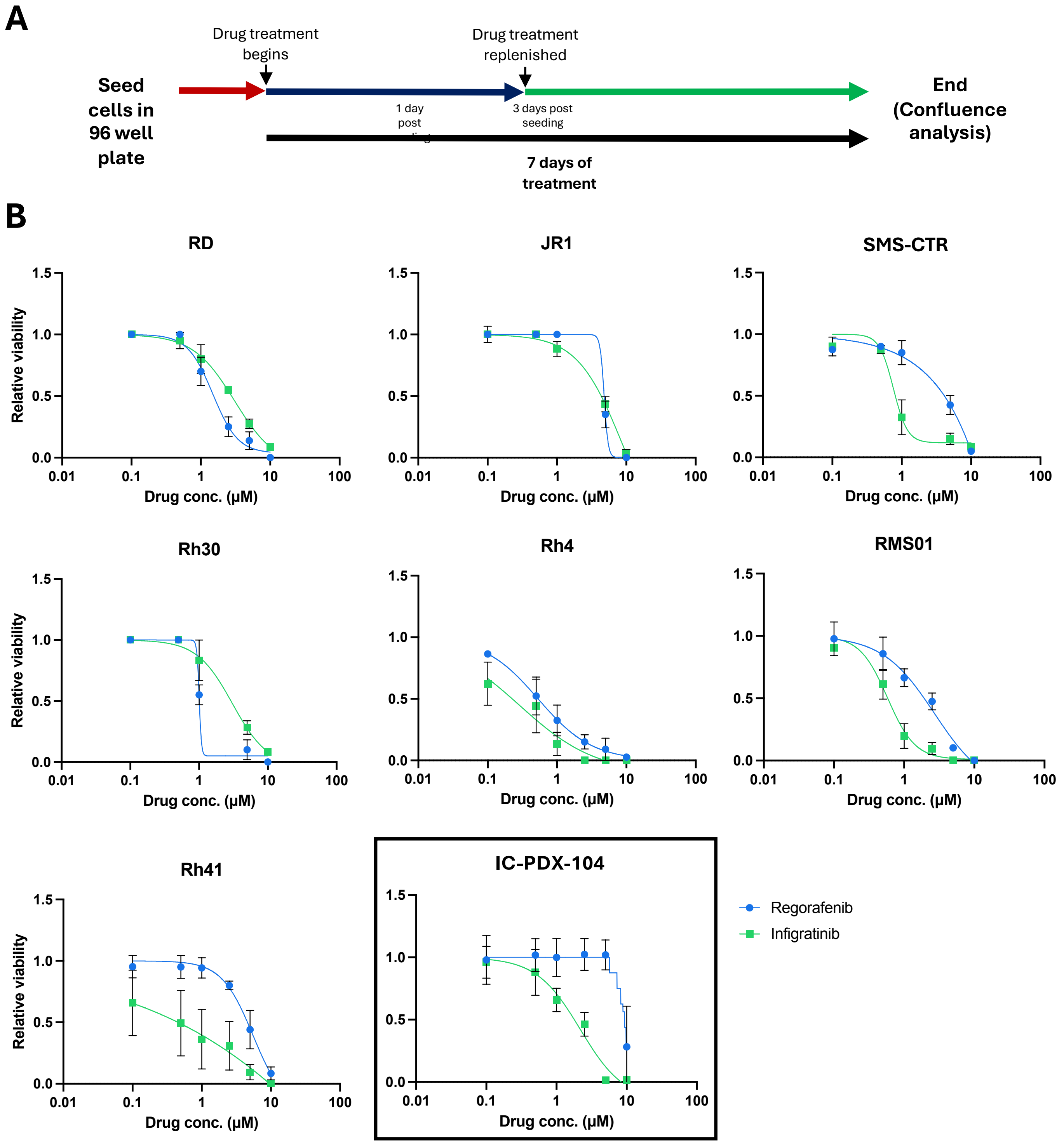

**Supplementary figure 6: Regorafenib and infigratinib treatment inhibits RMS *in vitro* cell line growth. A**, Schematic of the protocol for the RMS *in vitro* drug response assessment. **B**, non-linear fit dose response curves for regorafenib and infigratinib drug treatment in the 7 RMS cell lines and 1 patient derived culture. Data presented as mean values from three independent experiments +/- SEM.

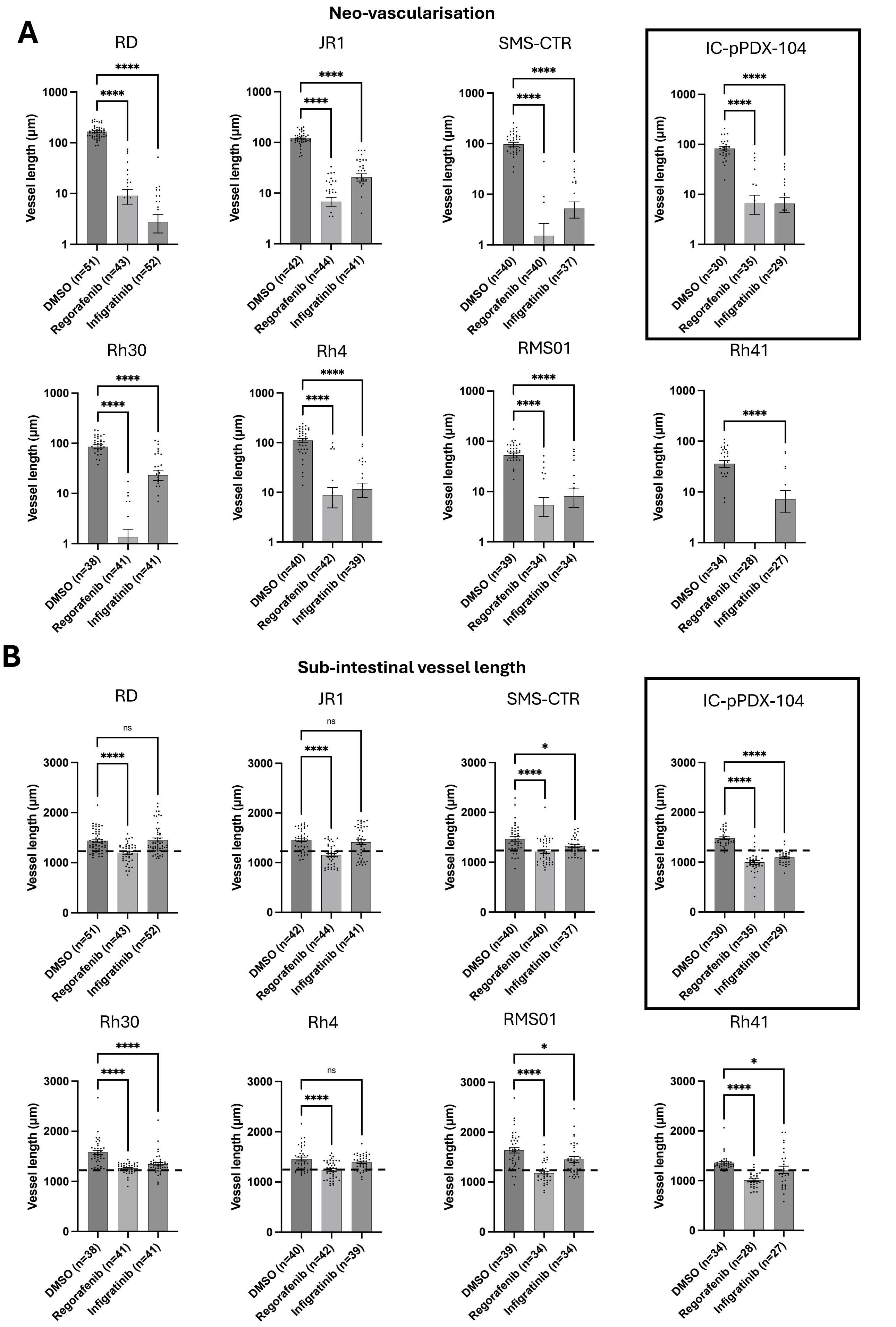

**Supplementary figure 7: Regorafenib and infigratinib treatment inhibits RMS xenograft tumour induced vascularisation.**

**Supplementary figure 7**: **Regorafenib and infigratinib treatment inhibits RMS xenograft tumour induced vascularisation.** Xenografts generated from the 7 RMS cell lines were treated with 0.1 μM regorafenib or infigratinib for 66 hours after which vessel development was assessed by FIJI image analysis. **A&B**, Bar graphs of absolute values of neo-vessel (**A**) and SIV length (**B**) for treated xenografts of each cell line. n = number of xenografts.

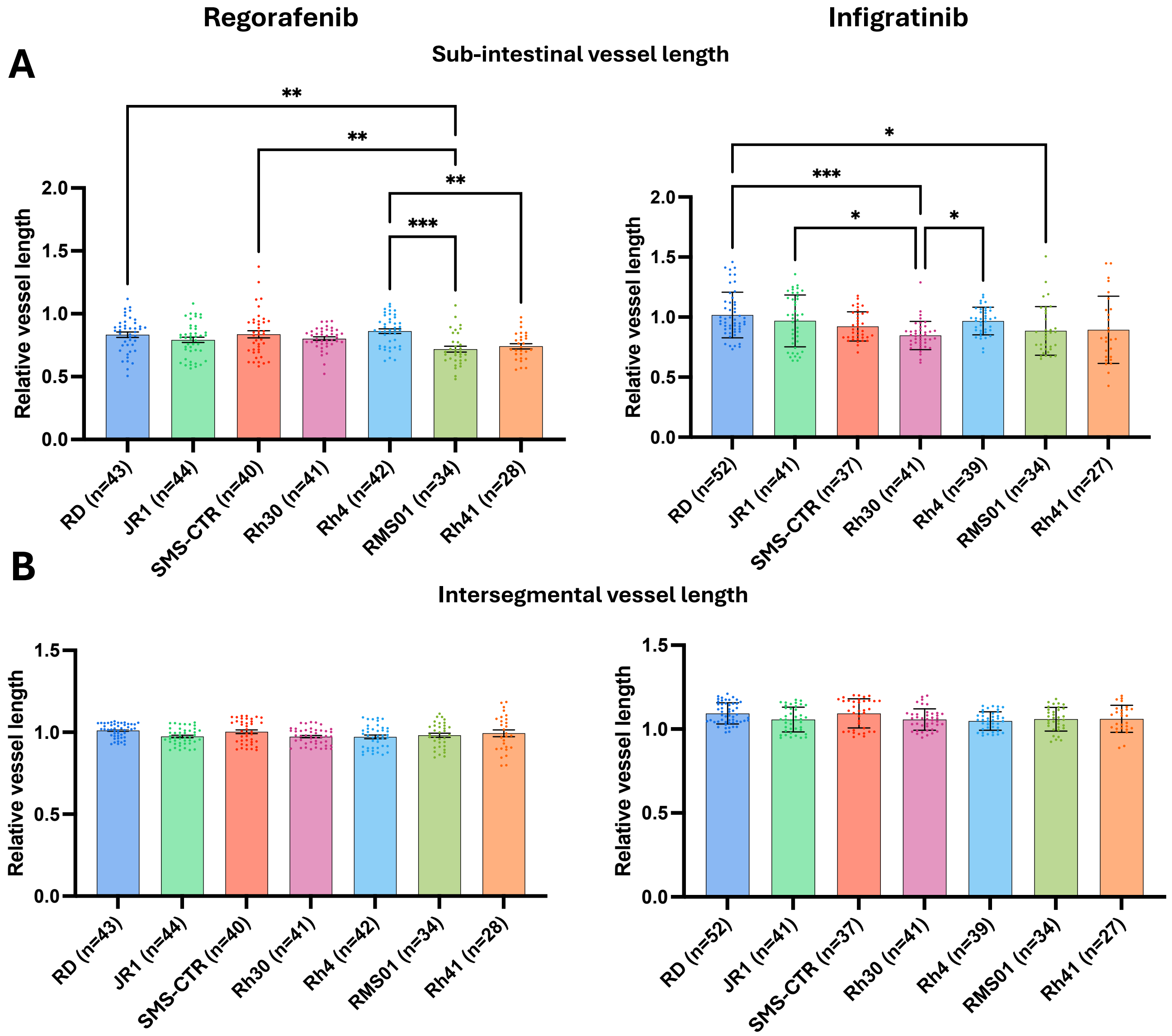

**Supplementary figure 8: Regorafenib and infigratinib treatment inhibits RMS xenograft tumour induced vascularisation.** Xenografts generated from the 7 RMS cell lines were treated with 0.1 µM regorafenib or infigratinib for 66 hours after which vessel development was assessed by FIJI image analysis. **A**, Bar graph of SIV lengths for treated xenografts relative to the average vehicle control treated vessel length for each cell line. **B**, bar graph of ISV length relative to the average vehicle control treated vessel length for each cell line. * P<0.05, ** P<0.01, *** P<0.001, **** P<0.0001, Ordinary one-way ANOVA with Tukey's multiple comparison's test, Data are presented as mean values +/- SEM. n = number of xenografts.

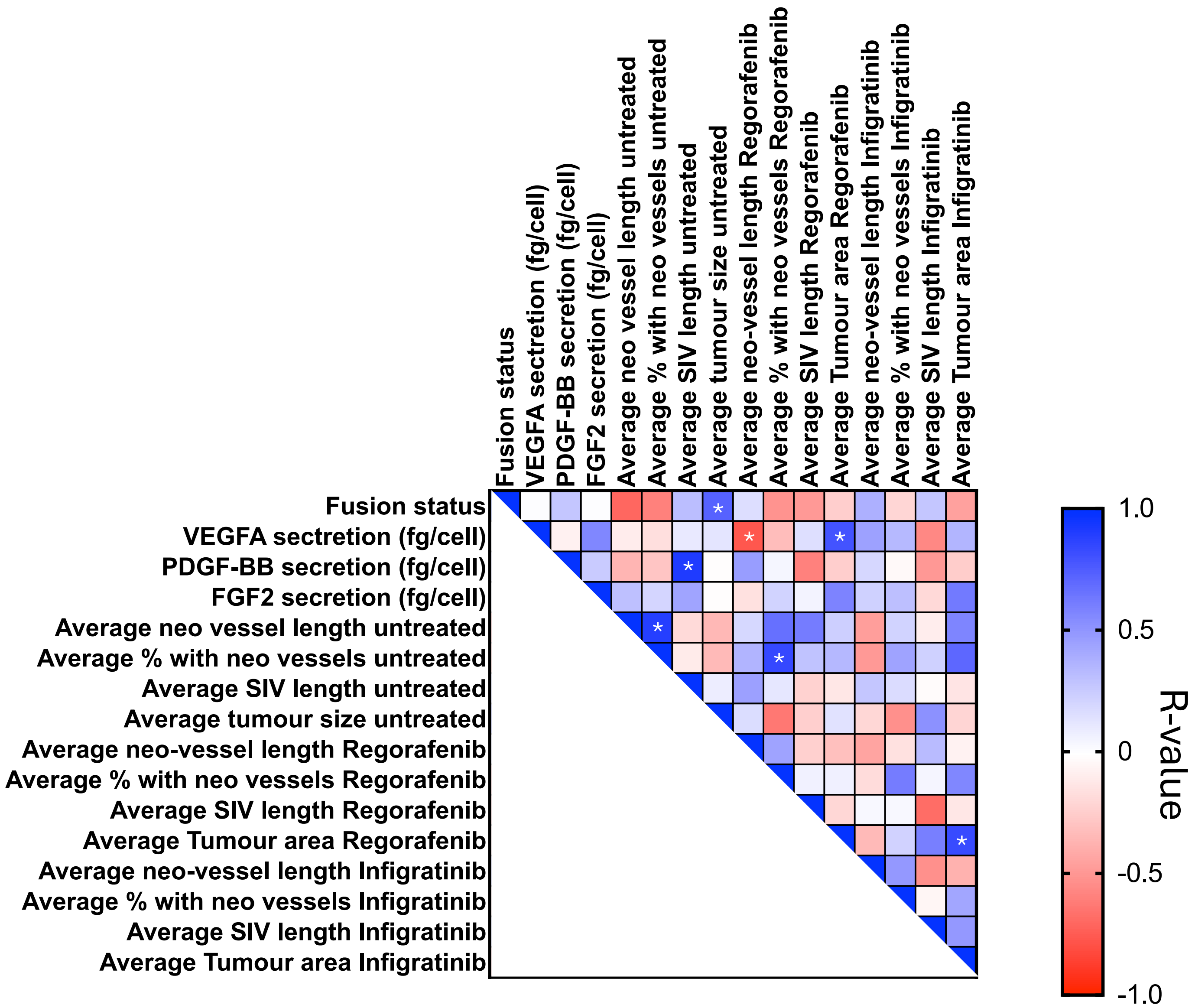

**Supplementary figure 9**: **Pearson correlation matrix** of Fusion status, angiogenic growth factor secretion, tumour and vessel growth metrics and response to MRTKIs regorafenib and infigratinib for all cell lines. R-values are shown. Asterix mark significant (P<0.05) associations (2-tailed).

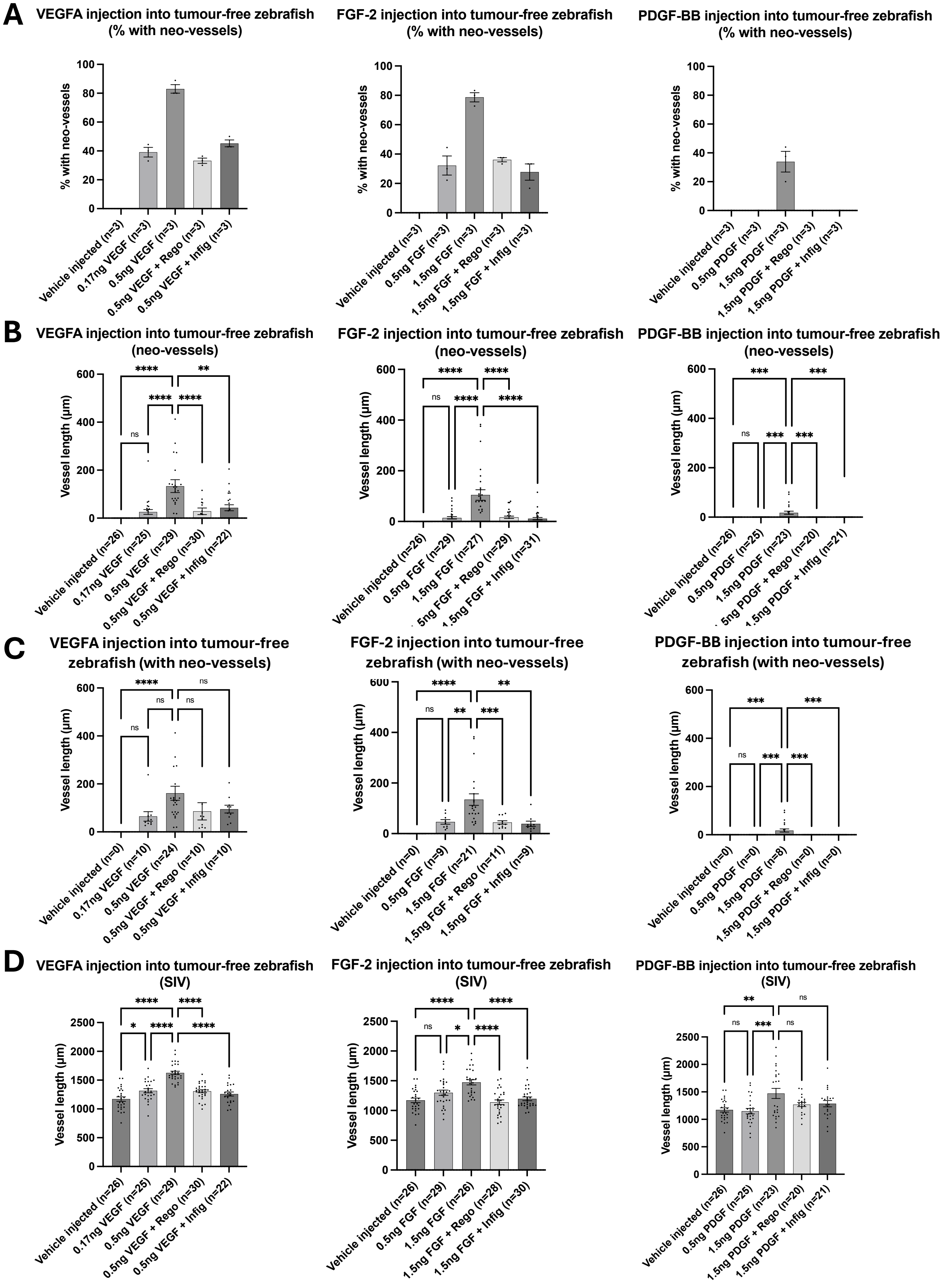

**Supplementary figure 10: Human recombinant VEGF, FGF-2 and PDGF-BB induce zebrafish vessel development which is blocked by Regorafenib and Infigratinib treatment.**

**Supplementary figure 10**: **Human recombinant VEGF, PDGF and FGF induce zebrafish vessel development which is blocked by regorafenib and infigratinib treatment.** 50 hpf *flk1:GFP* zebrafish embryos were injected with human recombinant VEGFA, FGF-2 and PDGF-BB to a final dose of 0.17-1.5ng. 54 hpf embryos were then treated with vehicle, 0.1 μM regorafenib or 0.1 μM infigratinib for 66 hours, prior to vessel imaging and analysis. **A-D**, Bar graphs showing the proportion of neo-vessel induction (**A**), average length of neo-vessels for all xenografts (**B**), average length of neo-vessels for xenografts where vessels were induced (**C**) and average SIV length (excluding neo-vessels) (**D**), with growth factor or vehicle control injection and drug treatment indicated (* P<0.05, ** P<0.01, *** P<0.001, **** P<0.0001), Ordinary one-way ANOVA with Tukey's multiple comparison's test, Data are presented as mean values +/- SEM). n = number of xenografts (except for A , which is number of experiments).
